# Breaking Barriers: Transitioning from X-ray Crystallography to Cryo-EM for Structural Studies

**DOI:** 10.1101/2025.10.13.682238

**Authors:** Hassan Zafar, Kiera L. Malone, Ajit K. Singh, Michael A. Cianfrocco, Karen C. Glass

## Abstract

Cryo-electron microscopy (cryo-EM) has transformed structural biology by enabling near-atomic resolution of large macromolecular complexes without the need for crystallization. Here, we describe our laboratory’s transition from X-ray crystallography to single-particle cryo-EM to investigate the ATPase family AAA+ domain-containing protein 2B (ATAD2B), a chromatin regulator implicated in epigenetic signaling. We outline the challenges encountered during protein expression, purification, and sample preparation, including co-purification of the chaperonin GroEL, and strategies employed to overcome these obstacles. Our workflow highlights critical steps in sample optimization, grid vitrification, and data processing using CryoSPARC, cisTEM, and Topaz, as well as computational requirements for high-resolution reconstructions. We also discuss model building, refinement, and validation approaches, emphasizing best practices for new cryo-EM users. This work provides practical insights for structural biologists adopting cryo-EM, particularly for large, flexible protein complexes, and underscores the importance of integrated approaches combining biochemical, computational, and imaging strategies.

## Background

### 1. The Expanding Toolkit of Structural Biology

The landscape of structural biology is constantly evolving, and significant advancements in cryo-electron microscopy (cryo-EM) methodologies have enabled us to study large macromolecular complexes of proteins and nucleic acids at near atomic resolution (Chua *et al*., 2022). Structural biology techniques such as X-ray crystallography, nuclear magnetic resonance, and electron microscopy have provided invaluable insights into the molecular machines that drive biological processes in the cell (Shoemaker & Ando, 2018). X-ray crystallography has long been the backbone of high-resolution structure determination, offering exquisite detail when well-diffracting crystals are available (Smyth & Martin, 2000, Zheng *et al*., 2015). However, the crystallization of many target biomolecules, particularly membrane proteins and large multicomponent complexes, has remained a challenge (Birch *et al*., 2018).

In cryo-EM, biological samples are applied onto specialized grids, which are then rapidly plunge-frozen (Thompson *et al*., 2016). Techniques like single-particle analysis have allowed researchers to achieve resolutions comparable to X-ray crystallography, while also revealing structural heterogeneities and dynamic intermediate states of molecular pathways (Cheng, 2015). However, the initial bottlenecks of employing cryo-EM include large investments in highly specialized electron microscopes and computational infrastructure (Meng *et al*., 2023). Additionally, the cryo-EM workflow, as illustrated in **Figure 1**, can be quite daunting.

**Figure 1.**
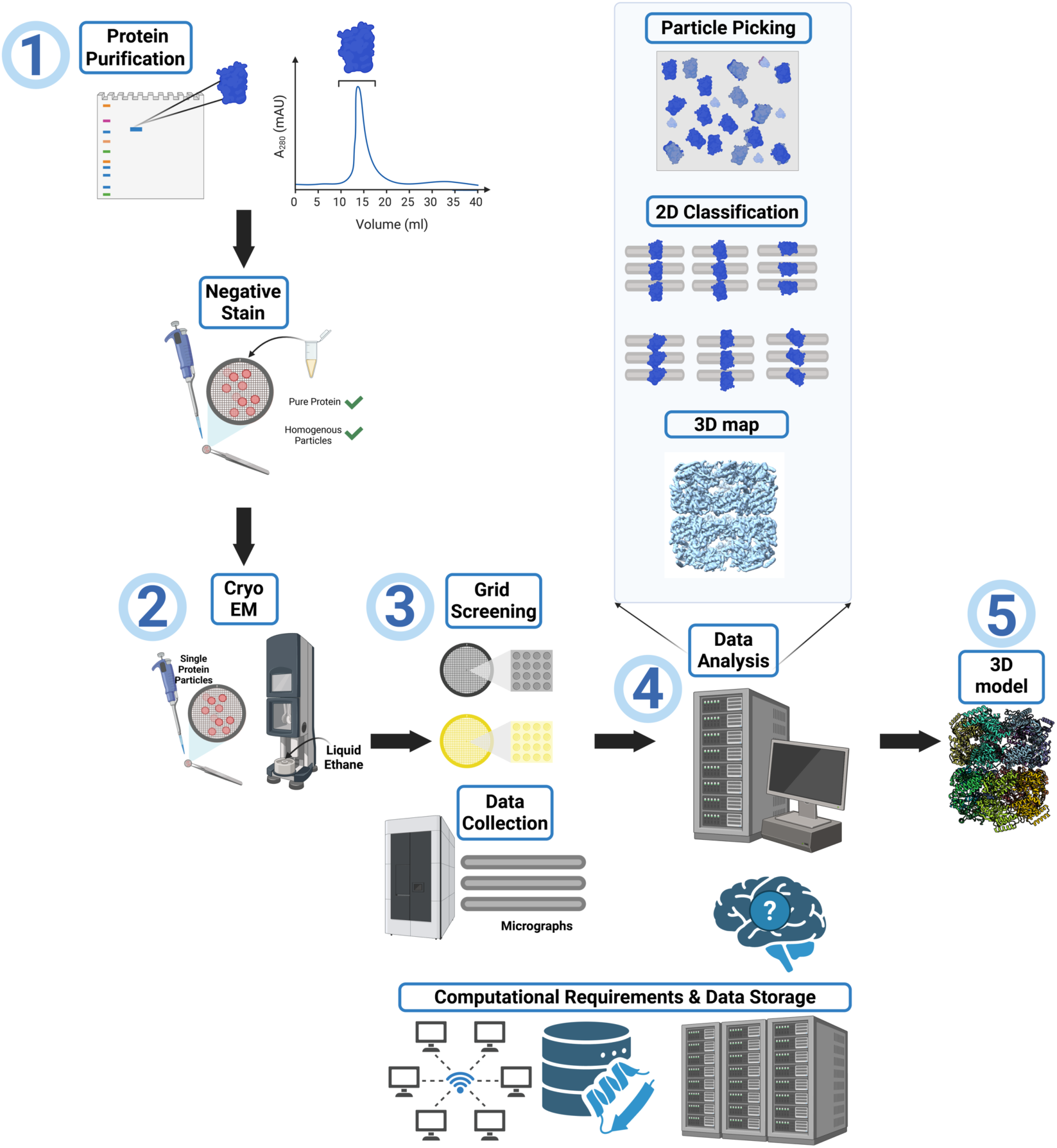
Overview of the cryo-EM workflow. Step 1: Evaluation of protein sample purity and quality to determine suitability for structural characterization by cryo-EM. Step 2: Vitrification of samples for cryo-EM imaging by applying the purified protein to a cryo-EM grid followed by plunge freezing. Step 3: Screening of vitrified cryo-EM grids to identify optimal freezing conditions for the purified sample. Parameters such as grid type, buffer components, additives, and plunge-freezing time/force are assessed. Once sample preparation is optimized, larger cryo-EM datasets are collected. Step 4: Data processing using specialized software, including motion correction, CTF estimation, particle picking, 2D classification, and 3D refinement, to obtain a high-resolution 3D map. Step 5: Model building and refinement to fit a model into the map. Model validation is performed to assess the quality of the map and the model fit. *Note: The cryo-EM workflow often requires iterative optimization at multiple stages, and users may need to revisit earlier steps to achieve a final structure at the desired resolution*.

Often, researchers must use an integrated approach, combining multiple techniques to obtain structural information on large biomolecular complexes (Nogales & Scheres, 2015). For instance, studies on large ribonucleoprotein complexes used the high-resolution information garnered with X-ray crystallography for individual subunits in conjunction with lower resolution cryo-EM maps of the multimeric complex to generate composite models (Bai *et al*., 2015). This synergy not only validates the individual findings of each method but also creates a more robust picture of the larger molecular architecture, which provides information that is essential for our understanding of biological mechanisms, and for the development of novel therapeutic interventions (Bijak *et al*., 2023).

### 2. Rationale for Developing Expertise in Cryo-EM for Structure Determination

#### Why Cryo-EM is Critical for Studying ATAD2B

In the Glass laboratory, we are interested in understanding how epigenetic signaling regulates gene expression and how alterations in these pathways are involved in disease development. Our research on bromodomain-containing proteins has focused on revealing the molecular mechanisms driving recognition of post-translational modifications found on histone proteins in the nucleosome. Bromodomains are a conserved structural motif that function as a chromatin reader domain to recognize acetylated lysine residues on histone proteins (Zeng & Zhou, 2002). Binding to specific acetyllysine modifications on histones often bridges the associated bromodomain-containing protein to chromatin where it can carry out its molecular activities. For the past several years our research has focused on the human ATPase family AAA+ domain-containing protein 2 (ATAD2), and its closely related paralog ATAD2B (Gay *et al*., 2019, Lloyd *et al*., 2020, Evans *et al*., 2021, Phillips, Malone*, et al.*, 2024). We have employed a variety of approaches including structural biology, molecular biology, genomics, biochemistry, biophysics, and proteomics to determine how physiologically abundant combinations of histone modifications regulate the reader activity of the ATAD2 and ATAD2B bromodomains. However, how the bromodomain region contributes to the cellular and molecular activities of ATAD2 and ATAD2B is unknown. These proteins are thought to function as regulators of chromatin architecture (Morozumi *et al*., 2016, Koo *et al*., 2016, Lazarchuk *et al*., 2020), and we hypothesized that recognition of acetyllysine residues in the flexible N-terminal region of histones plays an important role in directing the AAA+ ATPase activity of ATAD2B to its chromatin substrates. We quickly realized that using a structure-function approach to understand how ATAD2B contributes to these fundamental cellular mechanisms, designed to maintain chromatin organization, would necessitate isolating the full-length protein to study its activity *in vitro*.

The ATAD2B protein is 1,458 amino acid residues long and contains an unstructured N-terminal region, two AAA+ ATPase domains, a bromodomain, and an ordered C-terminal domain (**Figure 2a**) (Puchades *et al*., 2020, Khan *et al*., 2022). Previous studies on human ATAD2, and a yeast homolog Abo1, indicated that truncating the unstructured N-terminus would increase our ability to successfully express and purify the ATAD2B protein for biochemical and structural studies, without significantly impacting its enzymatic ATPase function (Cho *et al*., 2023, Cho *et al*., 2019). Thus, we codon optimized the DNA encoding the human ATAD2B protein residues 380-1458 for expression in *E. coli* and inserted it into the pGEX-6P-1 vector to add an N-terminal GST-affinity tag. We started with our well-established system in *E. coli* due to its ease of use and our previous experience purifying the ATAD2B bromodomain. However, the GST-ΔN-ATAD2B, res 380-1458 construct has a predicted molecular weight of 150 kDa, considerably larger than any other protein we purified previously (Singh *et al*., 2022, Poplawski *et al*., 2014, Phillips, Malone*, et al.*, 2024, Obi *et al*., 2020, Lubula, Eckenroth*, et al.*, 2014, Lubula, Poplawaski*, et al.*, 2014, Lloyd *et al*., 2020, Gay *et al*., 2019, Evans *et al*., 2021). For the remainder of the text, this GST-ΔN-ATAD2B construct (residues 380-1458) will be referred to as ATAD2B for clarity. Not surprisingly, the expression of ATAD2B in *E. coli* was very low, but after optimizing several factors including the IPTG concentration, buffer components, and the *E. coli* cell line we were able to reliably obtain ∼500 μL of 2.5 mg/mL ATAD2B from a four liter culture. However, there was a considerable amount of contaminating proteins remaining in the sample, even after the GST-affinity and size exclusion chromatography columns (**Figure 2b**).

**Figure 2.**
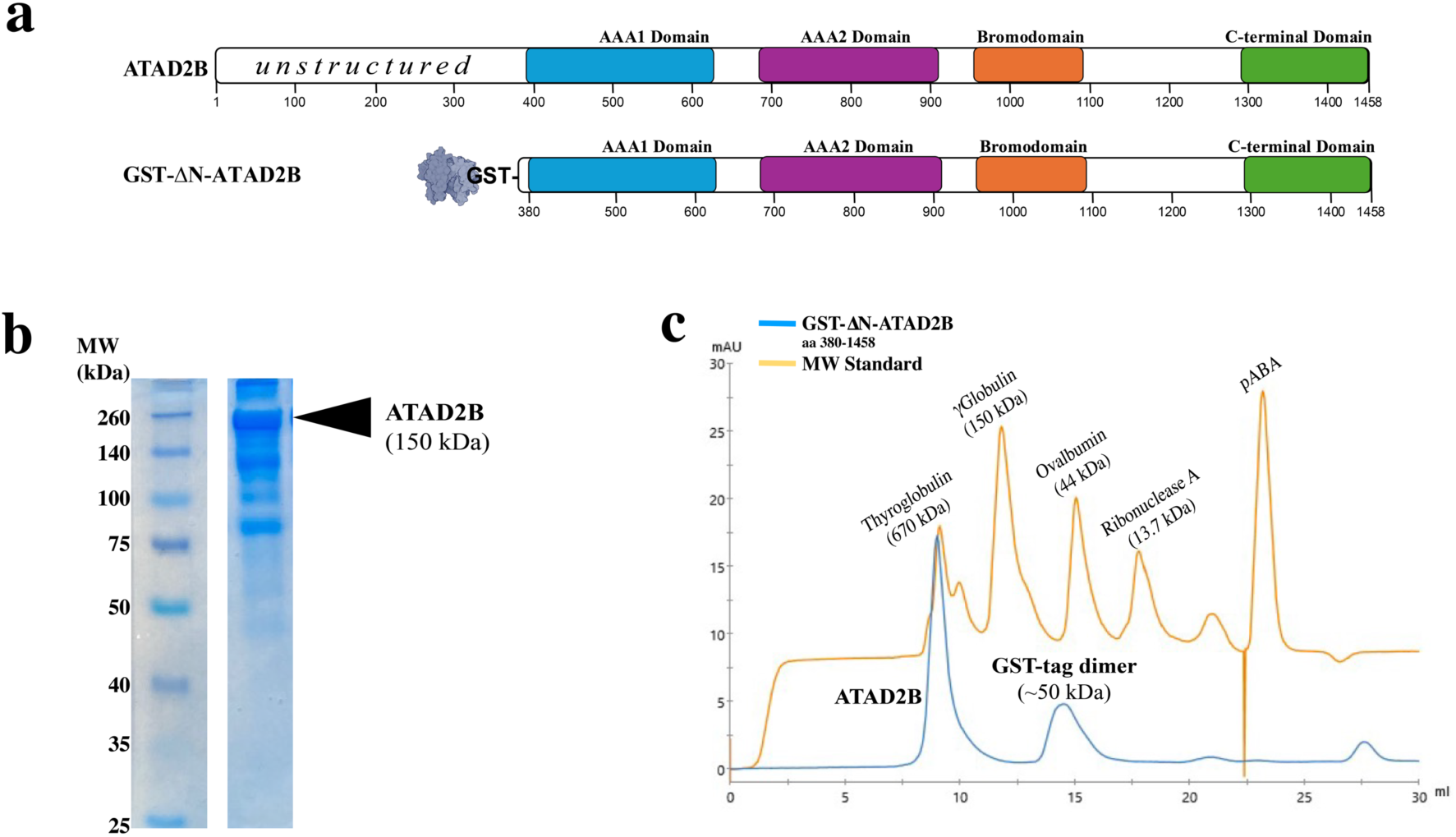
Evaluation of ATAD2B oligomerization. **a)** Domain organization of the full-length human ATAD2B protein (aa 1-1458), and the GST-tagged, N-terminally truncated-ATAD2B expression construct (aa 380-1458). For clarity, this GST-ΔN-ATAD2B construct is referred to as ATAD2B throughout the text. b) 10% SDS-PAGE gel stained with Coomassie blue showing the molecular weight (MW) standard and a lane corresponding to the size-exclusion chromatogram peak eluting at ∼ 9 mL (lane 2). In lane 2, ATAD2B appears as the upper band, migrating above its predicted molecular weight of 150 kDa. Additional bands in the same lane indicate sample heterogeneity. c) Size-exclusion chromatography traces from a Superdex 200 Increase 10/300 GL column (ÄktaPrime, GE/Cytiva) for the ATAD2B sample (blue) and a MW standard mixture (orange). Monomeric ATAD2B is expected at 150 kDa. The first peak (∼9 mL) elutes near thyroglobulin (670 kDa) consistent with ATAD2B forming an oligomeric complex. The second peak (∼15 mL), eluting before ovalbumin at (44 kDa), corresponds to dimerized GST-tags (∼50 kDa).

This sample was not pure enough, or in the milligram quantities needed for the X-ray crystallization approach we used previously with the ATAD2 and ATAD2B bromodomains (Phillips, Malone*, et al.*, 2024, Lubula, Eckenroth*, et al.*, 2014, Lubula, Poplawaski*, et al.*, 2014, Lloyd *et al*., 2020, Gay *et al*., 2019, Evans *et al*., 2021). Additionally, the 150 kDa ATAD2B AAA+ ATPase is expected to hexamerize into a ∼900 kDa complex upon ATP nucleotide binding (Puchades *et al*., 2020, Khan *et al*., 2022), and this size is well above the upper limit for structure determination using solution NMR, which is another structural determination approach we are familiar with (Poplawski *et al*., 2014, Kim *et al*., 2016, Obi *et al*., 2020, Evans *et al*., 2021, Singh *et al*., 2022, Phillips, Cook*, et al.*, 2024). The structure of the closely related yeast homolog of ATAD2B (Abo1) was determined using cryo-EM (Cho *et al*., 2019), which prompted us to begin learning more about this technique by reaching out to the National Center for Cryo-EM Access and Training (NCCAT) in New York, NY. We first heard about the NCCAT from attending structural biology meetings such as the American Crystallographic Association (ACA) and the Biophysical Society, along with word of mouth from colleagues. The NCCAT staff pointed us to several online cryo-EM resources (**Table 1**), including the Caltech Getting Started in cryo-EM course that includes lecture videos by Dr. Grant Jensen (Technology, n.d., Course, 2013). Fortuitously, we also attended the ACA annual meeting in Portland, OR, where we participated in a workshop on ‘Hands-on Single-Particle CryoEM Data Analysis with cryoEDU’ that was organized by Dr. Michael Cianfrocco. Additionally, a large portion of the scientific sessions at that meeting were focused on cryo-EM, including “New developments in cryo-EM”, “Machine learning in cryo-EM”, and “Structures of very large assemblies”.

**Table 1:**
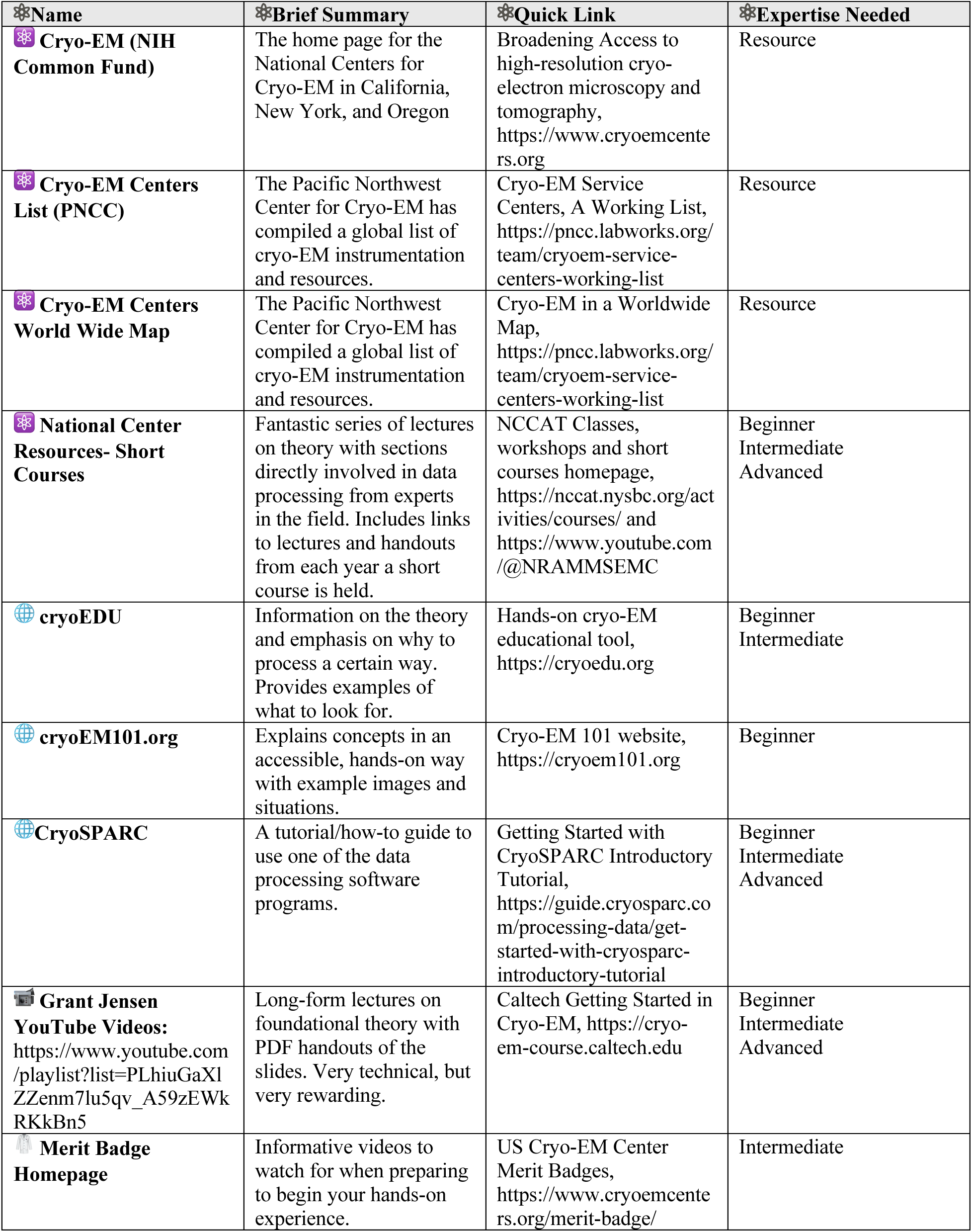
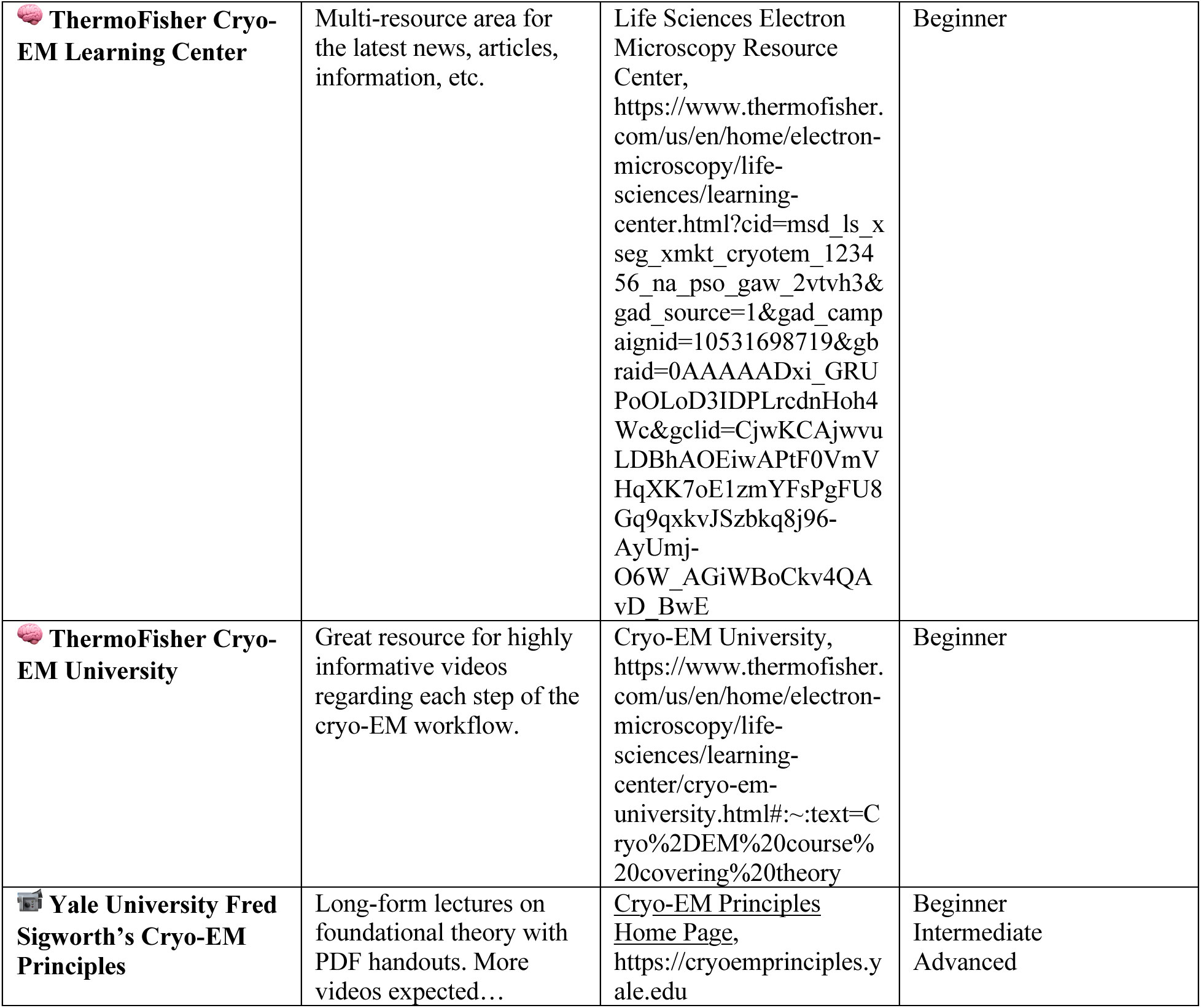
Helpful Resources for Getting Started in Cryo-EM.

The ACA meeting exposed us to the capabilities of cryo-EM, and we quickly discovered that cryo-EM requires significantly less protein than crystallography, that it can tolerate heterogenous samples, and that the ATAD2B AAA+ ATPase complex is an ideal size for structure determination using this method (Venien-Bryan *et al*., 2017). While structural biology techniques such as X-ray Crystallography and NMR require milligram quantities of highly purified protein samples to obtain high-quality/high-resolution data, it appeared to us that cryo-EM could handle sample heterogeneity a bit better (Takizawa *et al*., 2017, Wang & Wang, 2017), thanks to the downstream data processing workflow. Therefore, we decided it would be worthwhile to pursue learning cryo-EM to study the structure and function of human ATAD2B.

Early in this process, a postdoctoral trainee in the Glass laboratory had the opportunity to attend the cryo-EM short course at the NCCAT. As part of this course they were able to image the ATAD2B protein sample using negative stain microscopy (**Figure 1**, step 1), and the NCCAT staff also thought ATAD2B would be a good candidate to study using cryo-EM methods. To move forward we needed access to a plunge freezing system and a TEM microscope to make and screen cryo-EM grids. However, this expensive and specialized equipment was not available locally, so we applied to the TP1 training program at the NCCAT in NYC. To gain comprehensive, hands-on experience with every stage of cryo-EM sample preparation and data processing, four members of the Glass Lab traveled in pairs to the NCCAT for weeklong visits each month over the course of a year. When each team of two returned, they shared what was learned and their experiences troubleshooting with the rest of the lab. This allowed us to maximize our time with the NCCAT experts to learn the techniques necessary to determine the structure of the ATAD2B complex.

#### Helpful Resources for when you decide cryo-EM is right for you

For those interested in or new to cryo-EM, we highly recommend viewing introductory videos on cryo-EM theory and sample preparation. There are a plethora of different resources, and we have compiled many of them in **Table 1**. Additionally, virtual seminars, webinars, and scientific meetings are a great place to begin learning about cryo-EM. For those who want to learn more about cryo-EM facilities located nearby prior to investing in this technique, we would encourage readers to read the information available at each National Cryo-EM Center that are funded by the NIH-NIGMS. Moreover, Pacific Northwest cryo-EM Center (PNCC) has created a global list of available cryo-EM centers (see **Table 1**). We also encourage readers to visit the interactive map of ‘HighEnd CryoEM Worldwide’ that displays type of microscope, camera, and operation style, if appliable, in addition to contact information (**Table 1**). Reaching out to a facility directly will allow you to learn if the services they offer will fit your needs. For example, the national cryo-EM centers provide on-site training in addition to core services, whereas more regional institutions may accept submission of a purified protein sample, or a protein sample frozen on a grid.

### 3. Acquiring Technical Skills for Sample Preparation and Optimization

The transition from X-ray crystallography to cryo-EM involves modifications in sample preparation protocols, as the requirements for optimal imaging differ between the two techniques (Stark & Chari). In X-ray crystallography, the main goal is coaxing proteins or nucleic acids to form well-ordered crystals (Smyth & Martin, 2000). This requires optimization of various parameters such as pH, temperature, precipitating agents, and chemical additives—to discover the optimal conditions that promote crystal lattice formation. Well-ordered crystals with good diffraction yield high-resolution structural data (McPherson & Cudney, 2014).

For single-particle cryo-EM experiments, crystallization of samples is not required, though proteins and nucleic acids are often purified using conventional approaches (Saibil, 2022). Purified samples are applied to cryo-EM grids that feature a thick holey support film, typically made of carbon or gold, which provides a scaffold to support formation of vitreous ice. During plunge freezing, forming a uniform and thin layer of vitreous ice during plunge freezing is essential for capturing high-resolution images of macromolecules in a near-native state, andfor preserving the sample’s structural integrity under the electron beam(Passmore & Russo, 2016). In some cases, an additional ultra-thin layer of continuous carbon or graphene is added to directly support the biomolecules within the ice, helping to anchor them in place and maintain the integrity of fragile samples(de Martin Garrido *et al*., 2021). The choice of film material can influence particle distribution, orientation, and image contrast, and is often tailored to the specific requirements of the experiment (Lyumkis, 2019). Extensive optimization of the sample freezing conditions is often necessary to achieve uniform particle distribution and the desired ice thickness on the grid, both of which are critical for obtaining high-quality cryo-EM micrographs (**Table 2**) (Han *et al*., 2023). The quality of the cryo-grid sample directly impacts the success of downstream imaging and data analysis, thus making this step as critical in cryo-EM as crystallization is in X-ray studies.

**Table 2.** Factors to Consider During Cryo-EM Sample Preparation.

| Table 2. Factors to Consider During Cryo-EM Sample Preparation |  |
| --- | --- |
| Category | Condition / Instrument |
| Sample Type | Cryo-EM is ideal for large proteins and macromolecular complexes (e.g., over 50 kDa) that are difficult to crystallize. |
| Initial Screening | Acquire a small set of micrographs from negatively stained samples or frozen cryo-EM grids to evaluate particle integrity, homogeneity, and potential aggregation. |
| Buffer Optimization | Optimize salt (typically 150 -500 mM NaCl), glycerol (typically 0-5%), and pH (typically 6.5–8.5); consider adding: – Reducing agents (DTT, TCEP)– Detergents (e.g., OG) to reduce aggregation or preferred orientation– Cofactors or ligands (e.g., ATP, Mg<sup>2+</sup>) to stabilize functional complexes. |
| Protein Concentration | Test multiple concentrations to avoid aggregation (too high) or sparsity (too low). |
| Vitrification Devices | Compare methods (Vitrobot, Leica GP2, Chameleon) to minimize air–water interface damage. |
| Grid Type Selection | Use different grids (Quantifoil, UltrAuFoil, Graphene-coated) to optimize particle orientation and distribution. |
| Ice Quality Control | Maintain a clean, dry environment; use dried tools, LN<sub>2</sub> transfer boxes, and wear face masks to prevent frost and contamination. |

Transitioning to cryo-EM demands not only technical skill in operating transmission electron microscopes, but also familiarity with the specialized software used for data collection and analysis. Researchers must become adept at adjusting a wide range of settings—such as electron beam parameters, detector configurations, and imaging conditions—all of which play a crucial role in determining the quality of the resulting data. This involves understanding complex concepts such as defocus optimization, astigmatism correction, and dose management to minimize beam damage while maximizing signal quality. Additionally, practitioners must assess ice quality and particle distribution to optimize data collection strategies. This technical expertise represents a substantial investment in skill development that is essential for successful cryo-EM research. However, many national cryo-EM centers provide critical support to newcomers by offering guidance on selecting optimal experimental conditions, such as pixel size, total exposure, and target defocus, based on the size and characteristics of the protein under study. Facility staff also assist with assessing sample quality and advising on sample optimization, which is one of the primary motivations for establishing these centers. Such resources help ensure that researchers can achieve high-quality results even without extensive prior experience.

#### Gaining expertise in electron microscopy

Transitioning from X-ray crystallography to cryo-EM is like stepping into a new world, but one that speaks a familiar language. Our structural studies on ATAD2B began with crystallization attempts, which failed due to the inability to obtain high concentrations of purified ATAD2B. This challenge led us to pursue cryo-EM as an alternative approach. We initiated our efforts with negative staining to evaluate sample homogeneity, integrity, and concentration, key indicators for cryo-EM readiness using the JEOL 1400 TEM, 120 keV microscope, equipped with an AMT XR611 CCD camera available through the Microscopy Imaging Center at the University of Vermont Larner College of Medicine. As we lacked in-house vitrification tools, negative staining was the most useful way to initially evaluate sample quality. However, many facilities successfully produce cryo-EM structures without this step. When we brought the ATAD2B sample to the NCCAT for plunge-freezing we quickly realized that optimizing the sample conditions for cryo-EM grid preparation is as critical as optimizing buffer conditions for crystal growth. Variables such as salt, detergent, and glycerol can dramatically affect vitrification quality and particle distribution. High protein concentrations can lead to aggregation, while low concentrations result in sparse particle visibility. Vitrification of grids for cryo-EM introduces another level of complexity. Parameters like blotting time, blot force, temperature, and humidity all required optimization. Even the type of grid significantly influenced particle orientation and distribution (Weissenberger *et al*., 2021, Wang & Zimanyi, 2024). Recent advances in grid preparation strategies, including approaches to minimize air–water interface damage and improve particle distribution, are reviewed in Haynes et al. 2025 (Haynes *et al*., 2025). After testing multiple conditions, we found that a buffer containing 50 mM Tris pH 7.5, 150 mM NaCl, and 5% glycerol, vitrified on a 1.2/1.3 UltrAuFoil grid using a Vitrobot Mark IV, provided the best results in terms of particle orientation and distribution. Every step from ice thickness to grid clipping required precise, careful handling. These seemingly minor details were essential to enable successful high-resolution cryo-EM imaging.

#### Computational Requirements are an important factor for cryo-EM datasets

Cryo-EM data processing involves the handling of large datasets and requires substantial computational resources to perform key tasks such as motion correction, contrast transfer function (CTF) estimation, particle picking, 2D classification, and 3D refinement (Baldwin *et al*., 2018). For any lab entering the cryo-EM field, one of the first and most critical questions is which data processing suite(s) should you use and how do we meet the computational demands of these programs? When we began transitioning into cryo-EM, we faced the same challenge-should we invest in our own Graphics Processing Unit (GPU) workstations, rely on a cloud-based service, use a national shared facility, or work with the university’s high-performance computing (HPC) cluster? After evaluating our needs and budget constraints, we opted to use the Vermont Advanced Computing Center (VACC), which offered a cost-effective and scalable solution with routine data backups. To assist others in making informed decisions, **Table 3** summarizes the core computational requirements for commonly used software suites and platforms. These specifications represent minimum requirements for each program, and data processing performance can be significantly improved by upgrading hardware components such as GPUs, Random Access Memory (RAM), and file storage space.

**Table 3.** Core computation requirements for popular cryo-EM image processing platforms.

|  |  |  |  |
| --- | --- | --- | --- |
| Computational Requirements |  |  |  |
| Component (Minimum Requirements) | CryoSPARC (Punjani *et al.*, 2017) | RELION (Scheres, 2012) | cisTEM (Grant *et al.*, 2018) |
| CPU | ≥16 cores (Intel i9 / AMD Ryzen) | ≥16 cores (Intel i9 / AMD Ryzen) | ≥16 cores (Intel i9 / AMD Ryzen) |
| RAM | 64 GB | 64 GB | 64 GB |
| GPU | 1× NVIDIA RTX 3060 / A2000 | 1× NVIDIA RTX 3060 / A2000 | Not Supported |
| Fast Storage (SSD) | 1–2 TB NVMe SSD | 1–2 TB NVMe SSD | 1–2 TB NVMe SSD |
| Bulk Storage | ≥10 TB HDD or NAS | ≥10 TB HDD or NAS | ≥10 TB HDD or NAS |
| Network | 1 Gbps | 1 Gbps | 1Gbps |
| Operating System | Linux (Ubuntu, CentOS, RHEL) | Linux (Ubuntu, CentOS, RHEL) | Linux (Ubuntu, CentOS, RHEL) |
| Cryo-EM Platforms |  |  |  |
| Platform | Type | Notes | 🔗Quick Link |
| CryoSPARC Cloud | Cloud | Fully cloud-hosted by Structura Biotechnology Inc. |  |
| Open Science Grid (OSG) | Shared Facility HPC Cluster | National academic HPC resource | https://osg-htc.org |
| COSMIC<sup>2</sup> Platform | Shared Facility HPC Cluster | Centralized national resource | https://www.cosmic2.org |
| AWS Batch/EC2 | Cloud | Commercial cloud infrastructure | https://aws.amazon.com/products/ |
| ScipionCloud (Cuenca-Alba *et al.*, 2017) | Hybrid (Cloud + Shared/Local HPC) | Web interface; jobs can run on cloud or connect to local/university cluster |  |

### 4. Common Pitfalls and Problems

#### ATAD2B sample heterogeneity caused by a common contaminating protein

Early on in our data processing workflow, we observed heptameric particles in both our micrographs and 2D class averages. However, the cryo-EM structures of ATAD2 homologs were all hexameric (Cho *et al*., 2019, Cho *et al*., 2023, Wang *et al*., 2023). Initially, we hypothesized that these heptameric particles may support a new function for ATAD2B (Cho *et al*., 2019, Cho *et al*., 2023, Wang *et al*., 2023). To ensure that the heptameric particles were indeed ATAD2B, and not some unwanted protein, we sent both a liquid sample of ATAD2B and cutout bands from the SDS-PAGE gel of the ATAD2B sample shown in **Figure 2b** to a mass spectrometry core (also see supplemental data). In each sample, human ATAD2B was present. Because we were pushing the size limit for soluble protein expression in *E. coli* (Chae *et al*., 2017), these results suggested that the heterogeneity could be coming from sample degradation or internal cleavage of the ATAD2B protein. The aforementioned positive results gave us the confidence to continue with high resolution data collection. Our goal was to collect data on ATAD2B bound in three different nucleotide states to capture any large secondary/tertiary structural confirmational changes that occur between these states, and gain information on how nucleotide coordination changes within the binding pocket.

The NCCAT collected three Krios datasets: ATAD2B-apo (9,541 micrographs), ATAD2B-ADP (12,945 micrographs), and ATAD2B-γATP (10,880 micrographs)(**Figure 1**, step 3). Since ATAD2B is an ATPase, we used γATP, which is a commonly used slowly hydrolyzable ATP analog to “trap” ATAD2B in a nucleotide-bound state (Cho *et al*., 2019, Cho *et al*., 2023, Wang *et al*., 2023). The best data was obtained for the γATP dataset, and we started processing (**Figure 1**, step 4). After pre-processing, 2D classification, and a few rounds of refinement to remove junk, an initial heptameric 3D map to 3.3 Å was obtained. As our very first attempt at solving a cryo-EM structure, generation of this 3D map was an exciting milestone for our lab. We were eager to start building the ATAD2B protein model into this novel heptameric map (**Figure 1**, step 5). The predicted AlphaFold structure for the ATAD2B monomer (AF-Q9ULI0-F1-v4) was manually docked into the initial map using ChimeraX. However, despite trying multiple manual docking strategies, such as docking each domain separately into the map, making dimers, trimers, tetramers of the ATAD2B model for docking, and running fitinmap commands in ChimeraX, we were unable to fit any region of ATAD2B into the density map.

With progress in data processing halted, we decided it was time to reach out to a cryo-EM expert for help. We shared our 2D class averages and initial 3D model while explaining the docking issues, thinking that maybe our collaborator would have a better idea for fitting the ATAD2B model into the map (**Figure 3a**). The collaborator immediately identified our 2D class averages and 3D map as Gro-EL. Gro-EL is a ring-shaped *E. coli* chaperonin protein complex that helps mediate ATP-dependent polypeptide folding. It is an abundant, ubiquitous, and stress-induced protein that assists in folding proteins by interacting with incompletely folded polypeptides (Braig *et al*., 1994, Roh *et al*., 2017, Grimm *et al*., 1993, Fenton *et al*., 1994, Weissman *et al*., 1995). The presence of Gro-EL as a stress-induced protein made sense to us because we were pushing the upper limits of *E. coli* expression with the 150 kDa ATAD2B protein, which was expected to oligomerize into a 900 kDa complex(Khan *et al*., 2022). However, we were puzzled by the mass spectrometry results, which returned every single band cut out from the SDS-PAGE gel, and the liquid sample, as ATAD2B (see supplemental data). After reaching back out to the mass spectrometry core, we learned their analysis only searched our data against human protein databases. Once they opened up their search to include *E. coli* proteins, the analysis indicated the presence of Gro-EL in addition to ATAD2B.

**Figure 3.**
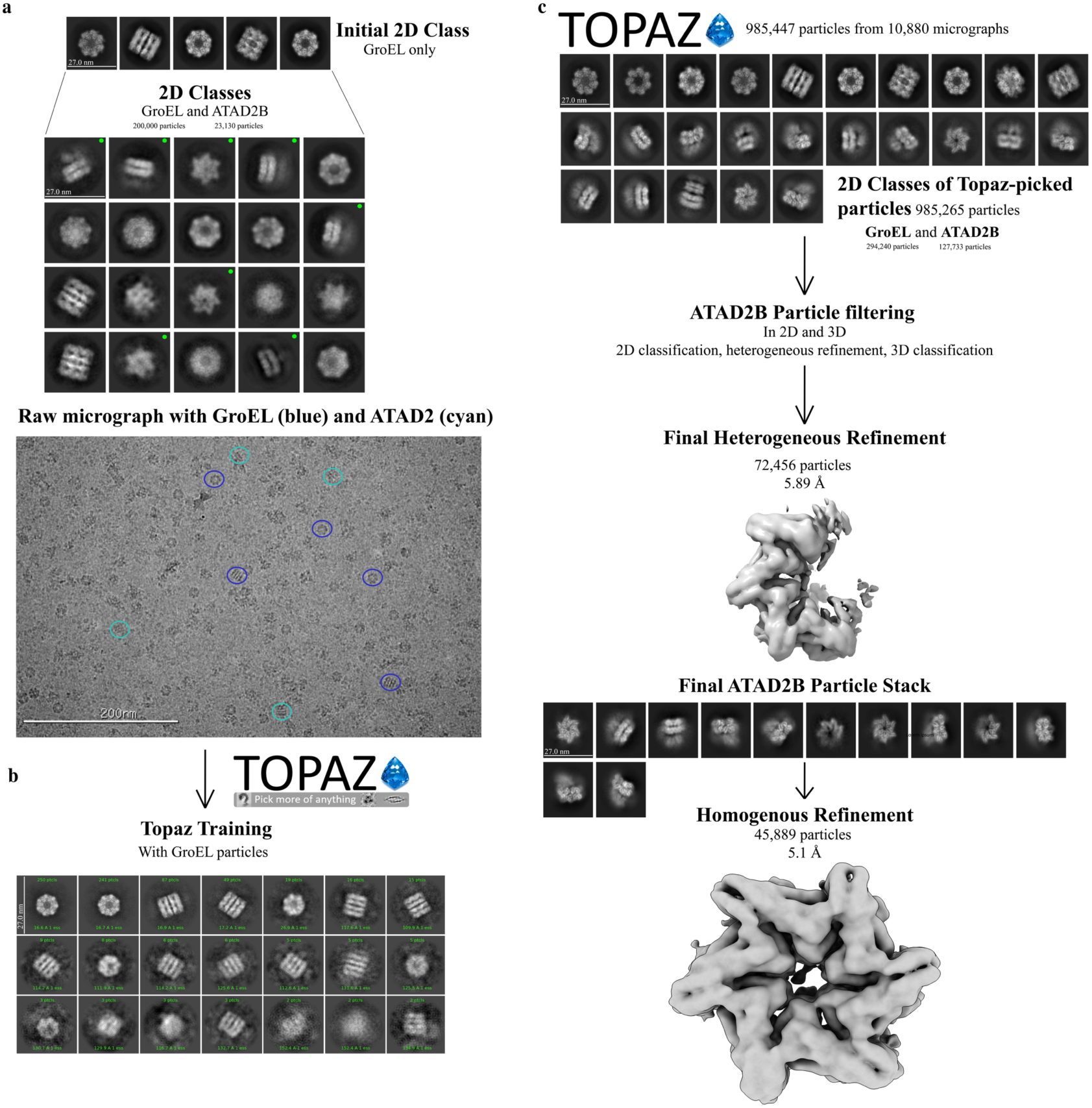
Cryo-EM data processing workflow for separating GroEL contaminants from ATAD2B particles. **a)** Initial 2D class averages from blob picking show predominantly heptameric GroEL particles (top). Further inspection reveals hexameric ATAD2B classes (marked with green dots, middle). A representative micrograph indicates the presence of both GroEL (blue circles) and ATAD2B particles (cyan circles). b) Topaz was trained to identify GroEL particles in raw micrographs. c) Topaz extract recovered more ATAD2B particles than expected. 2D class averages display GroEL (top row) and ATAD2B particles (middle and bottom rows) within the same dataset. Heterogenous refinement was used to separate the ATAD2B and GroEL particles, yielding a final ATAD2B map at 5.1 Å resolution from 45,889 particles.

Our analysis of the purified ATAD2B protein clearly indicate the sample is heterogeneous. A band is visible on the SDS-PAGE gel at the expected molecular weight for Gro-EL at 58 kDa (**Figure 4, top left panel**), which was confirmed via mass spectrometry. Gro-EL forms a stacked heptamer formed by 14 subunits, with a molecular weight of ∼812 kDa (Braig *et al*., 1994). The hexameric ATAD2B is a AAA+ ATPase is expected to have a molecular weight of ∼900 kDa. Therefore, these two large macromolecular complexes did not separate during analytical size-exclusion chromatography, and Gro-EL co-purified with ATAD2B throughout the entire process (**Figure 4, left panel**s). The SDS-PAGE gel and mass spectrometry data indicate that ATAD2B and Gro-EL exist in similar quantities in the sample (**Figure 4 top left, and supplementary data**). We had the unfortunate realization that our protein purification, sample preparation, and data collection strategies (**Figure 1**, steps 1-3) were optimized for imaging Gro-EL particles instead of ATAD2B. However, we did have a minor amount of ATAD2B particles visible in the micrographs, so the next step was to return to data processing (**Figure 1**, step 4) to determine if we could salvage our cryo-EM data and identify more ATAD2B particles.

**Figure 4.**
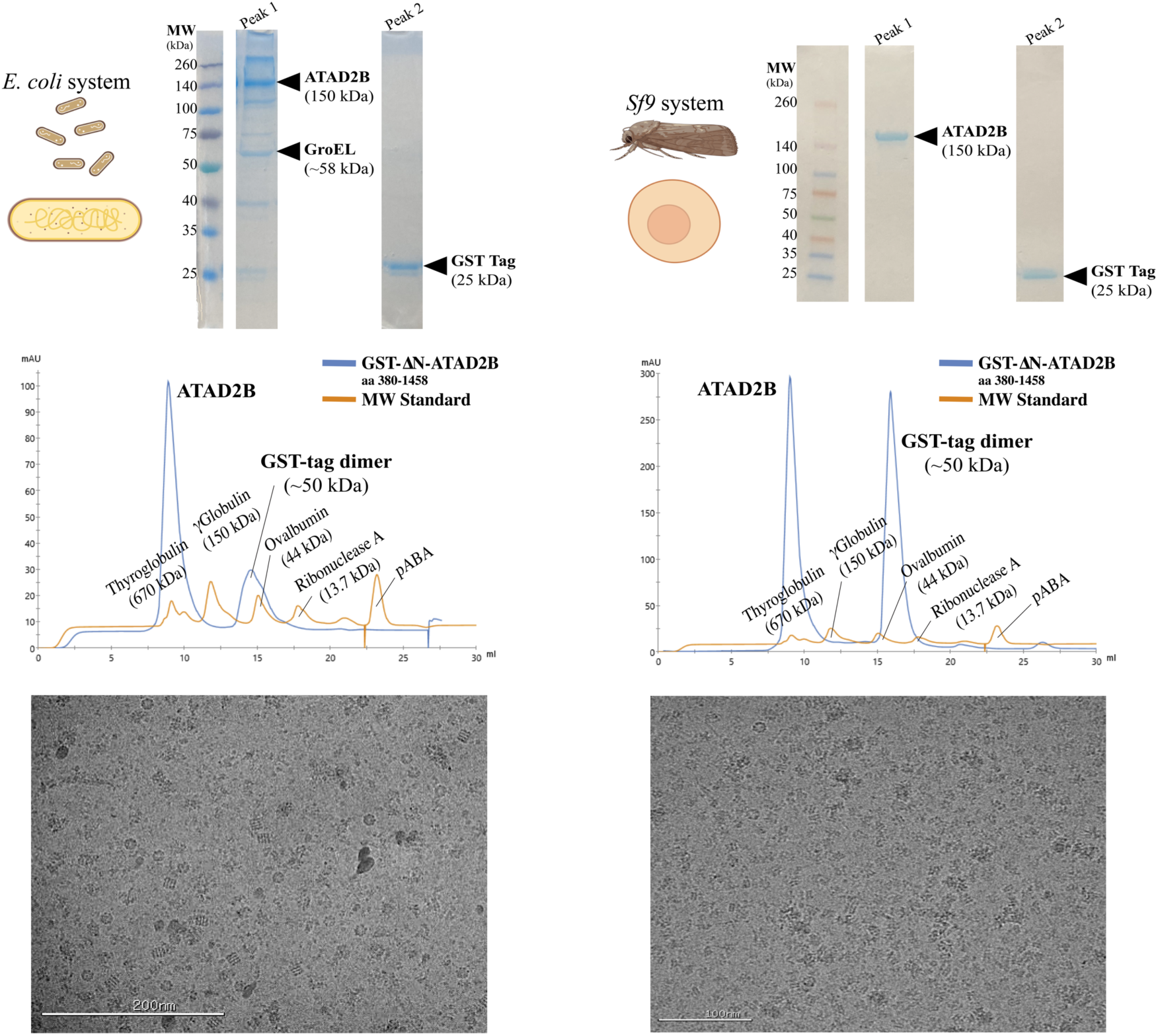
Comparison of ATAD2B expression and purification from *E. coli* and *Sf9* cells. Left panels: (Top) Schematic of *E. coli* expression Coomassie-stained 10% SDS-PAGE gel of peak samples from size-exclusion chromatography (SEC). The molecular weight (MW) ladder is shown in the first lane with the mass of each band labeled. Lane 2 shows protein sample from peak 1 of the SEC chromatogram below eluting at 9 mL that contains ATAD2B along with GroEL contaminant. Lane 3 shows protein sample from peak 2 eluting at ∼ 15 mL and contains the GST-tag. (Middle) SEC profile (Superdex 200 Increase 10/300 GL) showing a peak for ATAD2B and GroEL (9 mL). The additional peak at ∼15 mL corresponds to the cleaved GST-tag (observed as 50 kDa on SEC and 25 kDa on SDS-PAGE). (Bottom) Cryo-EM micrograph confirming the presence of heptameric GroEL contaminant in the purified sample. **Right panels**: (Top) Schematic of *Sf9* expression Coomassie-stained 10% SDS-PAGE gel showing greater than 90% pure ATAD2B collected from peak 1 of the SEC chromatogram below. The peak 2 lane contains the GST-tag (Middle) SEC profile (Superdex 200 Increase 10/300 GL) of the *Sf9*-purified sample with peaks for ATAD2B (∼9 mL) and the cleaved GST-tag (∼15 mLs). (Bottom) Cryo-EM micrograph confirming hexameric ATAD2B in the purified sample.

Before completely starting over in the data processing workflow, we reanalyzed the initial extracted stack of particles (**Figure 1**, step 4). After a few expansions of 2D class averages, additional top and side hexameric 2D class averages of ATAD2B were found (**Figure 3a**). In total, this particle stack contained ∼20,000 initial ATAD2B particles and ∼200,000 initial Gro-EL particles. To find more ATAD2B particles, we went back a step further to explore different particle picking strategies. Altering the settings for the standard blob picker and template picker jobs yielded no improvements in ATAD2B particle number. However, we had the good fortune to attend a seminar on Topaz, which is a neural network based particle picking software that gets “trained” on a small subset of particles known to be your protein of interest, to then find these particles in the full dataset (Bepler *et al*., 2019). We started to use Topaz with our ATAD2B particle stacks, but because of how training works, it was nearly impossible to get an accurate picking model. To avoid introducing noise or false picks for ATAD2B, the higher quality and easier to find Gro-EL particles were used in the Topaz training step. We also decided it would be helpful to continue learning cryo-EM data processing with the Gro-EL particles, so we could apply this workflow later, once we had better quality data on ATAD2B. Incredibly, training Topaz on the Gro-EL particles not only picked Gro-EL particles, but also picked more ATAD2B particles than any other strategy tried thus far (**Figure 3b**). Topaz found 127,733 initial ATAD2B particles and 294,420 initial Gro-EL particles, indicating we had many ATAD2B particles in this sample despite the Gro-EL contamination. However, after filtering to remove junk particles from the Topaz picks, only <100,000 ATAD2B particles remained to use in reconstructions. Unfortunately, there were not enough ATAD2B particles present generate a map beyond mid-resolution (∼5 Å) that would be needed to observe the details of nucleotide coordination in the ATAD2B binding pocket (**Figure 3c**).

We realized that it was time to return to the biochemistry (**Figure 1**, step 1) to obtain a more homogenous protein sample for cryo-EM structural investigation, without Gro-EL. There are a few protocols available to remove the Gro-EL contamination from a protein sample. One includes addition of unfolded bacterial lysate supplemented with ATP to try and outcompete Gro-EL from your protein of interest (Rohman & Harrison-Lavoie, 2000). We tried this purification method with a few rounds of optimization, however, Gro-EL still appeared at the same intensity on SDS-PAGE gels after purification. The final test was another cryo-EM screen and small overnight dataset collection, which confirmed that Gro-EL was present in the same amount prior to the addition of the unfolded bacterial lysate.

We had to decide if we should switch protein expression systems or collect enough cryo-EM data with the Gro-EL contaminant to obtain a higher resolution structure. In the γATP dataset, 10,880 micrographs were collected and less than 200,000 initial ATAD2B particles were observed before all of the junk was removed. Taking these numbers into consideration, we decided that it would require too much microscope time to make it worthwhile to continue with this sample. It was time to switch expression systems and return, again, to the biochemistry (**Figure 1**, step 1). In the literature, cryo-EM structures of ATAD2 homologs Abo1 and Yta7 were successfully determined from protein samples generated with the Sf9 insect cell expression system (Cho *et al*., 2019, Cho *et al*., 2023, Wang *et al*., 2023). We formed a new collaboration with Trybus Lab at the University of Vermont, who generously taught us how to express the ATAD2B protein using Sf9 inset cells. Within four months we obtained pure ATAD2B protein, free of Gro-EL, suitable for downstream cryo-EM studies (**Figure 4**). Obtaining a high-quality protein sample was essential for determining the structure of ATAD2B at near-atomic resolution, enabling us to characterize ligand coordination and protein–protein interactions that underpin its mechanism of action. This requirement parallels X-ray crystallography, where the quality of the crystal dictates the clarity of the diffraction data. Similarly, in cryo-EM, the purity and stability of the protein govern the resolution and reliability of the final structure—underscoring that sample preparation is the foundation for revealing atomic-level details in both techniques.

#### Protein expression with chaperone and other common contaminants

The presence of chaperones and other contaminants are a frequent challenge in cryo-EM sample preparation that complicate the data collection and analysis pipeline **(Table 4)** (Zielinski *et al*., 2021). Molecular chaperones such as the chaperonin GroEL can co-purify with target proteins, as they bind unfolded or partially folded polypeptides, making them unwanted entities during purification (Weissman *et al*., 1995, Rutledge *et al*., 2022). Expression and purification of the AAA+ ATPase ATAD2B complex in *E. coli* resulted in GroEL contamination that co-purified with our target protein. Due to the similar sizes of both proteins, we did not realize the chaperone was present in our data until the 2D classification stage in data processing. Our attempts to separate the GroEL protein from ATAD2B during purification were unsuccessful. Therefore, we reverted to using insect cell lines of *Spodoptera frugiperda* (*Sf9*) instead of bacterial protein expression systems for protein expression and purification. Moreover, based on our experiences it can be safely stated that effective cryo-EM studies require careful purification strategies, such as additional chromatography steps and nuclease treatments, as well as vigilant screening of grids to identify and minimize these potential contaminants.

**Table 4.** Common Cryo-EM Contaminants.

| **Table 4. Common Cryo-EM Contaminants** |  |
| --- | --- |
| **Protein contaminant** | **Notes** |
| **ArnA, SlyD and AcrB** (Andersen *et al.*, 2013, Caliseki *et al.*, 2025) | Frequently contaminate samples with His-tagged fusion proteins expressed in *E. coli* due to their natural histidine-rich sequences. |
| **GroEL / GroES** (Nain *et al.*, 2022) | Very common chaperonin contaminant from *E. coli* expression systems. |
| **Ferritin** (Wenborn *et al.*, 2015) | Typically encountered in samples derived from animal tissues or produced using mammalian expression systems. |
| **Ribosomal proteins / fragments** (Bolanos-Garcia & Davies, 2006, Singh *et al.*, 2020, Amunts *et al.*, 2014) | Can be found in protein samples purified from bacterial or mammalian cells. |
| **Heat shock proteins (Hsp70, Hsp90, sHsps)**(Carr *et al.*, 2025, Bolanos-Garcia & Davies, 2006) | These chaperones often bind unfolded proteins and co-purify with them. |
| **Affinity tag fragments (Glutathione S-transferase (GST) / Maltose-binding protein (MBP)** (Waugh, 2011) | Tag remnants are common contaminants in proteins purified using affinity tags. |

To help future labs convert to this new technique and avoid spending a lot of time troubleshooting removal of a contaminating protein, we would like to pass on what was helpful to us during this process. In **Table 5**, we have included a list of specialized data processing programs. The most common cryo-EM data processing software or platforms are found in **Table 3**, while a collection of helpful resources for learning various aspects of cryo-EM methods are mentioned in **Table 1**. We have included a list of common contaminants in **Table 4** to help assess micrographs for their presence. We also encourage readers to investigate specific contaminants and why they may occur. Knowing when to switch expression systems will vary between labs. We chose to switch due to the amount of microscope time that would be required to collect a high-quality dataset. We needed to apply for time at the National CryoEM Center, which is a limited resource, and we felt that it would be better to create a better biochemical sample to collect a high quality dataset. Despite a higher upfront investment in time to switch expression systems, we decided moving to Sf9 cells would set the lab up future studies on large complexes that interact with chromatin. For more information on undesirable contaminations, and other issues that may impact image quality, such as ice thickness, contamination, preferred orientation, etc., we refer you to these reviews ((Tan *et al*., 2017),(Cianfrocco & Kellogg, 2020),(Neselu *et al*., 2023), and (Wang & Zimanyi, 2024)), and to Chapter 3 at https://cryoem101.org/chapter-3/ for guidance.

**Table 5.** Specialized Modular Programs for Data Processing.

| Table 5. Specialized Modular Programs for Data Processing |  |  |
| --- | --- | --- |
| *Name | *Brief Description | *Quick Link |
| **AlphaFold**<br>(Jumper *et al.*, 2021) | AI system for predicting protein 3D structures from amino acid sequences. | https://alphafold.ebi.ac.uk/ |
| **ChimeraX**<br>(Pettersen *et al.*, 2021) | Molecular visualization program for displaying and analyzing 3D biological structures, including atomic models and cryo-EM maps | https://www.cgl.ucsf.edu/chimerax/ |
| **CryoDRGN**<br>(Zhong *et al.*, 2021) | Deep learning for heterogeneous cryo-EM data. | https://cryodrgn.csail.mit.edu/ |
| **crYOLO**<br>(Wagner *et al.*, 2019) | Neural network-based particle picker for cryo-EM micrographs. | https://cryolo.readthedocs.io/en/stable/ |
| **DynaMight**<br>(Schwab *et al.*, 2024) | A tool to model continuous conformational changes in cryo-EM datasets | https://github.com/3dem/DynaMight/ |
| **EMAN2**<br>(Tang *et al.*, 2007) | Suite for single-particle reconstruction and electron microscopy image processing. | https://blake.bcm.edu/emanwiki/EMAN2 |
| **Phenix**<br>(Adams *et al.*, 2010, Afonine *et al.*, 2018, Liebschner *et al.*, 2019) | Software suite for macromolecular structure determination by X-ray crystallography and cryo-EM. | https://phenix-online.org/ |
| **ModelAngelo**<br>(Jamali *et al.*, 2024) | Automated atomic model building for cryo-EM maps. | https://github.com/3dem/model-angelo/ |
| **Topaz**<br>(Bepler *et al.*, 2019) | Machine learning particle picking for cryo-EM. | https://cb.csail.mit.edu/topaz/ |

### 5. Data Processing Software for Single-Particle Cryo-EM Workflows

Data processing for single-particle cryo-EM is a multi-step, computationally intensive workflow that requires specialized software and robust infrastructure. Each stage, ranging from motion correction and CTF estimation to particle classification and high-resolution refinement, demands significant computing power and storage. Commonly used software suites such as CryoSPARC (Punjani *et al*., 2017), cisTEM (Grant *et al*., 2018, Lucas *et al*., 2021), and RELION (Scheres, 2012, Kimanius *et al*., 2016, Zivanov *et al*., 2022) (summarized in **Table 3**) often operate in tandem, generating terabytes of data from raw movies to final reconstructions. These requirements necessitate high-performance computing (HPC) clusters equipped with powerful GPUs, multi-terabyte storage, and high-speed networking. At the University of Vermont, collaboration with VACC staff was critical for building and maintaining this infrastructure, enabling parallel processing and efficient large-scale data transfers. Given the size of cryo-EM datasets, effective data management strategies are critical, and should include long-term storage plans for older datasets because downstream analyses may require access to raw data. Emerging trends such as cloud-based solutions and AI-driven tools further highlight the evolving nature of cryo-EM data processing.

#### Tutorial on data processing using the E. coli GroEL datasets to generate map reconstructions

The GroEL containing samples, expressed and purified from *E. coli,* were processed for cryo-EM analysis. The initial GroEL-γATP dataset was analyzed using CryoSPARC v.4.5.31 (Punjani *et al*., 2017). All 15,169 movies underwent pre-processing, including patch-based motion correction. Patch contrast transfer function (CTF) estimation was done on the motion-corrected micrographs. This is essential for high-resolution reconstructions, as it enables more precise correction of image distortions. The two parameters of ice thickness and CTF fit were checked for the selection of 14,568 micrographs for further processing. The next step included picking particles from the micrographs. In CryoSPARC there are different methods to do this such as manual picking, blob picking, template and deep learning methods like Topaz (Bepler *et al*., 2019). The circular blob picker in CryoSPARC identifies particles in micrographs by detecting circular features of a specified size. It provides a fast, template-free method for initial particle selection. We used blob picker to pick 10,124,313 particles. The initially picked particles were filtered using defocus-adjusted power and pick scores, which help assess the quality and reliability of each pick. These metrics allow for the exclusion of low-quality or spurious particles before extraction. The selected particles were then extracted from the micrographs using a box size of 300 pixels, ensuring that each particle is fully captured for downstream processing. A stack of 3,056,751 particles was subjected to reference free 2D classification. This type of classification groups particles into a set number of classes by aligning and averaging them in-plane, improving the signal-to-noise ratio of each class average. This makes it easier to identify and remove poor-quality or contaminant classes. A total of 317,080 particles were used to generate six *ab-initio* classes. Three of the six *ab-initio* classes, comprising at total of 262,461 particles, resembled GroEL and were selected to generate a homogeneous reconstruction using D7 symmetry. A homogeneous map with D7 symmetry is a three-dimensional reconstruction in which all selected particles are refined together under the constraint of seven-fold dihedral symmetry, which can enhance resolution by leveraging the inherent symmetry of the complex. GroEL is known to exhibit D7 symmetry, as confirmed by previously published studies (Torino *et al*., 2023). To further analyze structural heterogeneity, the selected particles were divided into four distinct 3D classes without applying a focused mask, allowing for unbiased classification across the entire volume. This approach facilitates the separation of different conformational states and the removal of heterogeneous or junk particles. One of the resulting classes was identified as junk and excluded from further processing. The other three classes comprising 261,106 particles were combined to generate a final non-uniform map of 3.3 Å, according to the gold standard 0.143 FSC cutoff criteria. A scheme for the data processing is shown in **Figure 5**.

**Figure 5.**
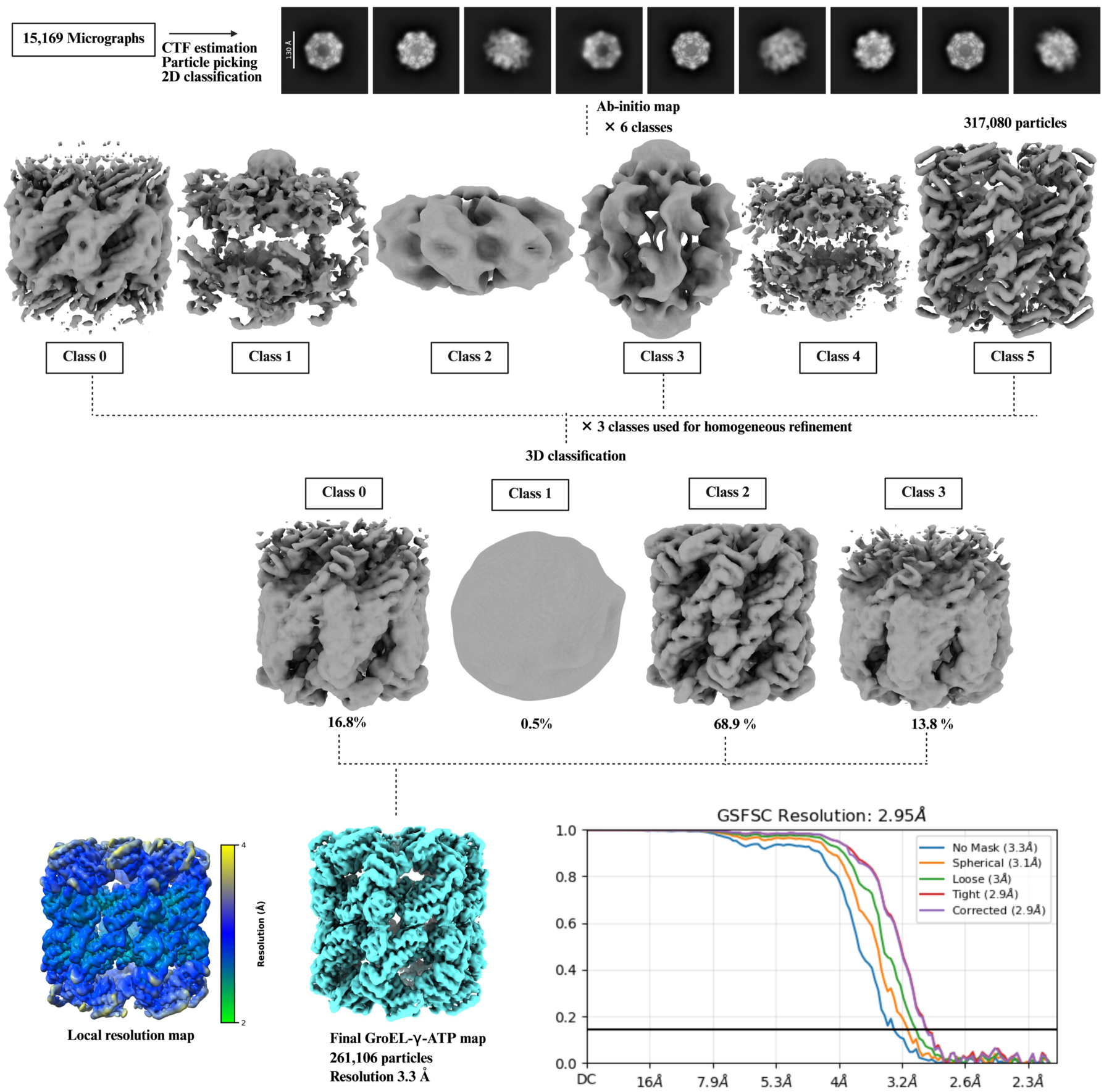
Single-particle cryo-EM data processing scheme for the GroEL-γATP bound complex. A total of 317,080 GroEL particles were selected following 2D classification and subjected to *ab initio* reconstruction into six classes. Three of these *ab initio* classes were retained for further processing and used to generate a homogeneous 3D map. Subsequent 3D classification yielded four classes, with one class identified as junk and excluded from further analysis. The remaining three classes were combined, resulting in a final reconstruction comprising 261,106 particles with a CryoSPARC gold standard FSC resolution of 2.95 Å, however, we report the no mask resolution of 3.3 Å for the final map. Local resolution estimation was performed in CryoSPARC, and the resulting map was visualized using UCSF ChimeraX.

The GroEL-apo dataset comprising 9,541 micrographs was also processed using CryoSPARC v.4.5.31 (Punjani *et al*., 2017). Patch contrast transfer function (CTF) estimation was carried out on the motion-corrected micrographs. The processing scheme was similar to the GroEL-γATP dataset. However, in this dataset we first performed manual picking from selected micrographs to create a high-quality training set. This manually curated set of particles was then used to train a Topaz model (Bepler *et al*., 2019), enabling more accurate automated particle picking for the entire dataset. A total of 513,428 particles were extracted from the micrographs with a box size of 256 pixels. From the 2D classification, a total of 148,146 particles were used to generate one *ab-initio* class. This class was used to generate a homogeneous map with D7 symmetry and was further classified into 5 classes. The final map generated from non-uniform refinement had a resolution of 3.7 Å according to the 0.143 FSC cutoff criteria. The processing scheme is shown in **Figure 6**.

**Figure 6.**
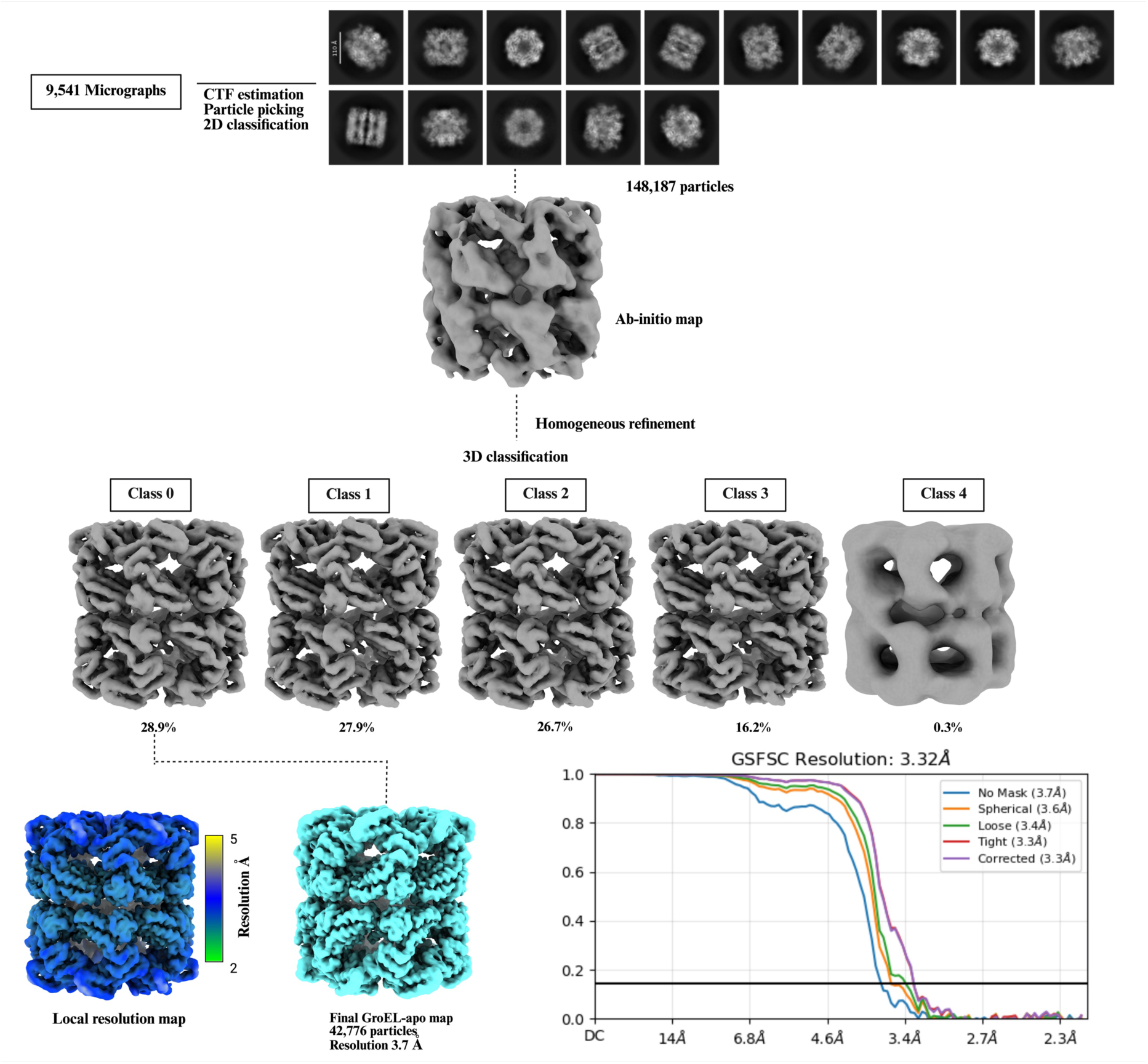
Single-particle cryo-EM data processing scheme for apo GroEL complex. A total of 148,187 GroEL particles were selected after 2D classification and subjected to *ab initio* reconstruction. Homogeneous refinement was performed on the resulting single class. Subsequent 3D classification yielded five classes; adopting a conservative approach, only the highest-quality class was selected for final map generation. The final reconstruction comprised 42,776 particles and had a final resolution of 3.32 Å according to the CryoSPARC gold standard FSC. However, we report our final map as 3.7 A with no mask applied. Local resolution estimation was performed in cryoSPARC, and the resulting map was visualized using UCSF ChimeraX.

For the GroEL-ADP dataset we chose to use a different software called cisTEM (Grant *et al*., 2018) for data processing. 11,46,722 particles were picked using the *ab-initio* template picker a were subjected to reference free 2D classification. Particles in the 2D classes resembling GroEL were selected, and a stack of 158,542 particles was exported into FREALIGN (Grigorieff, 2016). The particle stack was binned 8× 4×, 2× using the command resample.exe. This binning of the particle stacks is done to reduce the computational processing time. To prevent the introduction of high resolution bias a low pass filtered (20 Å) map of GroEL was generated from the PDBID: 9C0C, and was used for initial particle alignment. The 2× binned stack was then 3D classified into 8 classes. The classes with the largest number of particles were combined to generate a final 1X map at a resolution of 4.2 Å. The processing scheme is shown in **Figure 7**.

**Figure 7.**
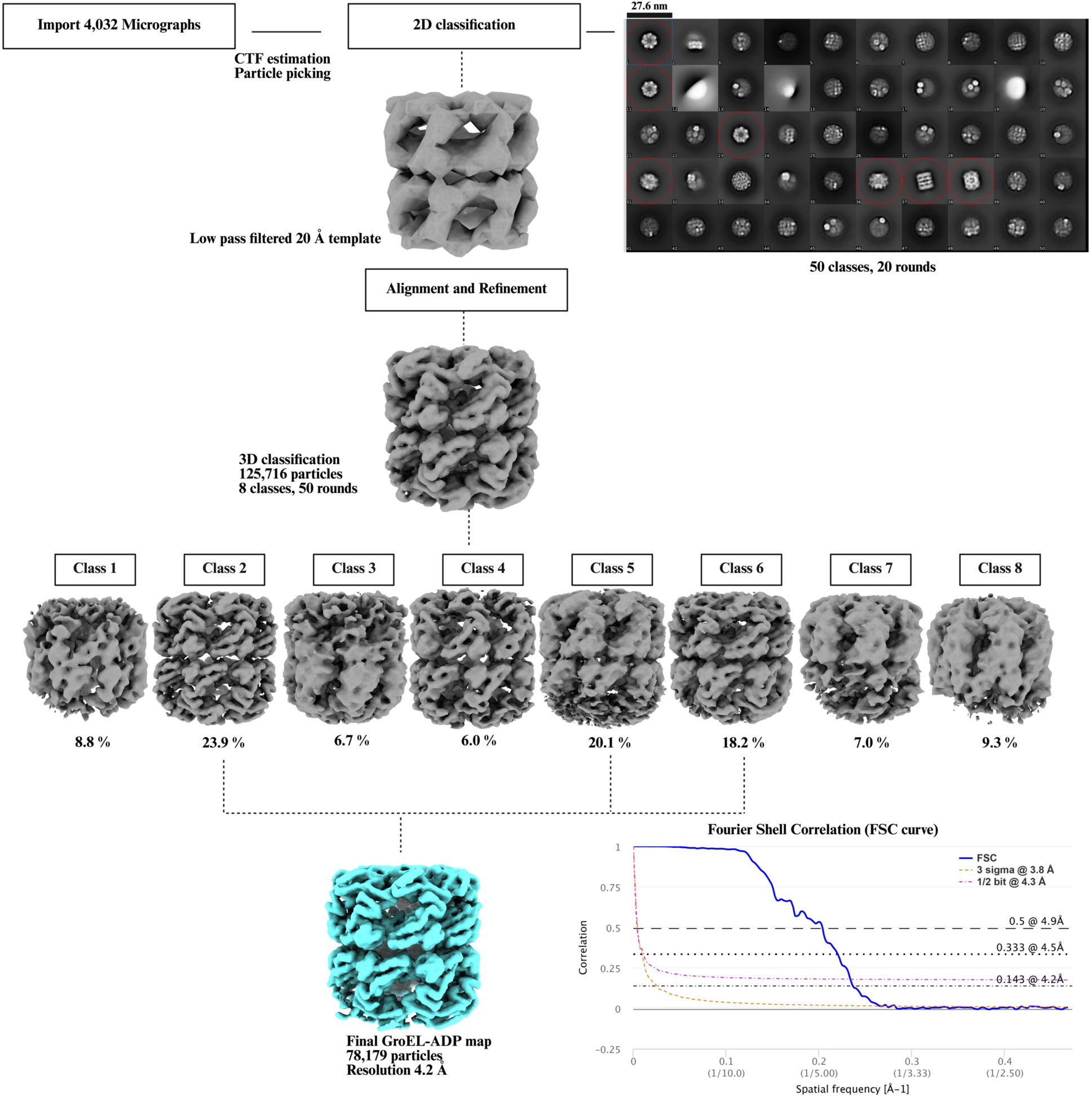
Single-particle cryo-EM data processing scheme for the GroEL-ADP bound complexes. The particle stack, comprising 125,716 GroEL particles selected from a 2D classification in cisTEM, was exported for further processing in FREALIGN. An initial alignment was performed using a low pass filtered GroEL template. Subsequently, the aligned particles underwent 3D classification over 50 rounds to sort them into distinct conformational states. However, no major confirmational changes were observed in the 3 best classes. These classes were combined to generate a final map of 4.2 Å. The two half-maps generated from the final 3D reconstruction were used to compute a Fourier Shell Correlation (FSC) curve by using the EMDB validation tool (https://www.ebi.ac.uk/emdb/validation/fsc/).

### 6. Nuances of Model Building, Refinement, Cryo-EM Structure Validation and Deposition

Once the highest-quality reconstruction has been achieved, the next step is to interpret the cryo-EM density map by building an atomic model. This model may be derived from previously solved structures obtained through techniques such as X-ray crystallography or cryo-EM, or predicted using computational tools like AlphaFold. Regardless of the source, model building should be guided by prior biological knowledge and experimental context to ensure accurate interpretation. An integrated approach is essential, often combining structural, biochemical, and biophysical data to inform model construction and validation. Techniques such as mass spectrometry, cross-linking mass spectrometry (XL-MS), small-angle X-ray scattering (SAXS), nuclear magnetic resonance (NMR), size-exclusion chromatography (SEC), and SEC coupled with multi-angle light scattering (SEC-MALS) can provide valuable insights into the composition, organization, and conformational states of the molecular complex.

Macromolecular crystallography and cryo-EM differ fundamentally in how structural information is obtained. In X-ray crystallography, diffraction patterns obtained from highly ordered crystals are used to compute electron density maps, which provide atomic-level detail when crystals diffract well. A major challenge is the “phase problem”. While diffraction experiments measure the intensities of scattered X-rays, they do not capture the corresponding phase information, which is essential for calculating an accurate electron density map. Because phases cannot be obtained directly from the experiment, specialized experimental strategies or computational methods must be used to estimate or recover them in order to solve crystal structures (Adams *et al*., 2009).

In contrast, cryo-EM reconstructs three-dimensional Coulomb potential maps from many thousands of particle images of flash-frozen molecules captured in random orientations. Unlike X-ray crystallography, cryo-EM micrographs retain the phase information, avoiding the crystallographic phase problem. While cryo-EM maps often exhibit lower signal-to-noise ratios and variable resolution across flexible regions (Punjani & Fleet, 2023), high-resolution areas can be interpreted with reduced model bias. Model building in cryo-EM typically involves docking known protein fragments or predicted structures into the density, followed by iterative refinement to improve the fit and identify secondary structure elements (Venien-Bryan *et al*., 2017).

#### Model building of GroEL in different nucleotide bound states

The initial steps of model building for X-ray crystallography and cryo-EM differ significantly due to the nature of the data (Terwilliger *et al*., 2020). However, after the initial fitting of the starting model in the final map (**Figure 1**, step 5), these two techniques become similar. For our three GroEL structures, we used PDBIDs 8BL7 and 9C0C as starting models. Here, we will focus on the model building of GroEL-γATP, scheme shown in **Figure 5**. The initial model was rigid body fitted in the final map using UCSF ChimeraX (Pettersen *et al*., 2004). As the name indicates, it is just placing the initial rigid model in the map based on optimal position and coordinates without changes to the model’s shape. The coordinates for the γATP ligand (AGS) were taken from PDBID: 5DAC and was rigid body fitted into the map. Further manual adjustments were done using Coot (Crystallographic Object-Oriented Toolkit) (Emsley & Cowtan, 2004), which is a molecular graphics program for building and validating macromolecular models. Our GroEL model was then structurally refined in the cryo-EM map using phenix.real_space_refine in Phenix (Liebschner *et al*., 2019). This program refines the atomic coordinates while simultaneously enforcing good stereochemistry, such as ideal bond lengths, angles, and proper protein backbone and side-chain conformations.

During the refinement steps, secondary restraints files were generated for the model and were used in subsequent refinement cycles. The restraints prevent the model from deviating from physically and chemically plausible conformations. Without these restraints, the refinement software would be at risk of overfitting the model to noise or artifacts in the cryo-EM map, leading to a geometrically unreliable structure. During refinement in Phenix, the resolution-dependent model-to-map correlation was evaluated using the correlation coefficient plots generated at the end of the process. These plots confirmed that all three final atomic models exhibited high correlation with their respective experimental density maps, indicating an excellent fit without evidence of overfitting. Separately, validation of the resulting models indicated favorable stereochemical parameters, including minimal deviation from ideal bond lengths and angles, suggesting high quality throughout the model geometry. Structural details of GroEL-γATP are also shown in **Figure 8**, with closer views of the nucleotide binding site of chain A. Figures were generated using UCSF ChimeraX (Pettersen *et al*., 2004) and PyMOL (version 2.3.1, Schrödinger, LLC) (DeLano, 2002).

**Figure 8.**
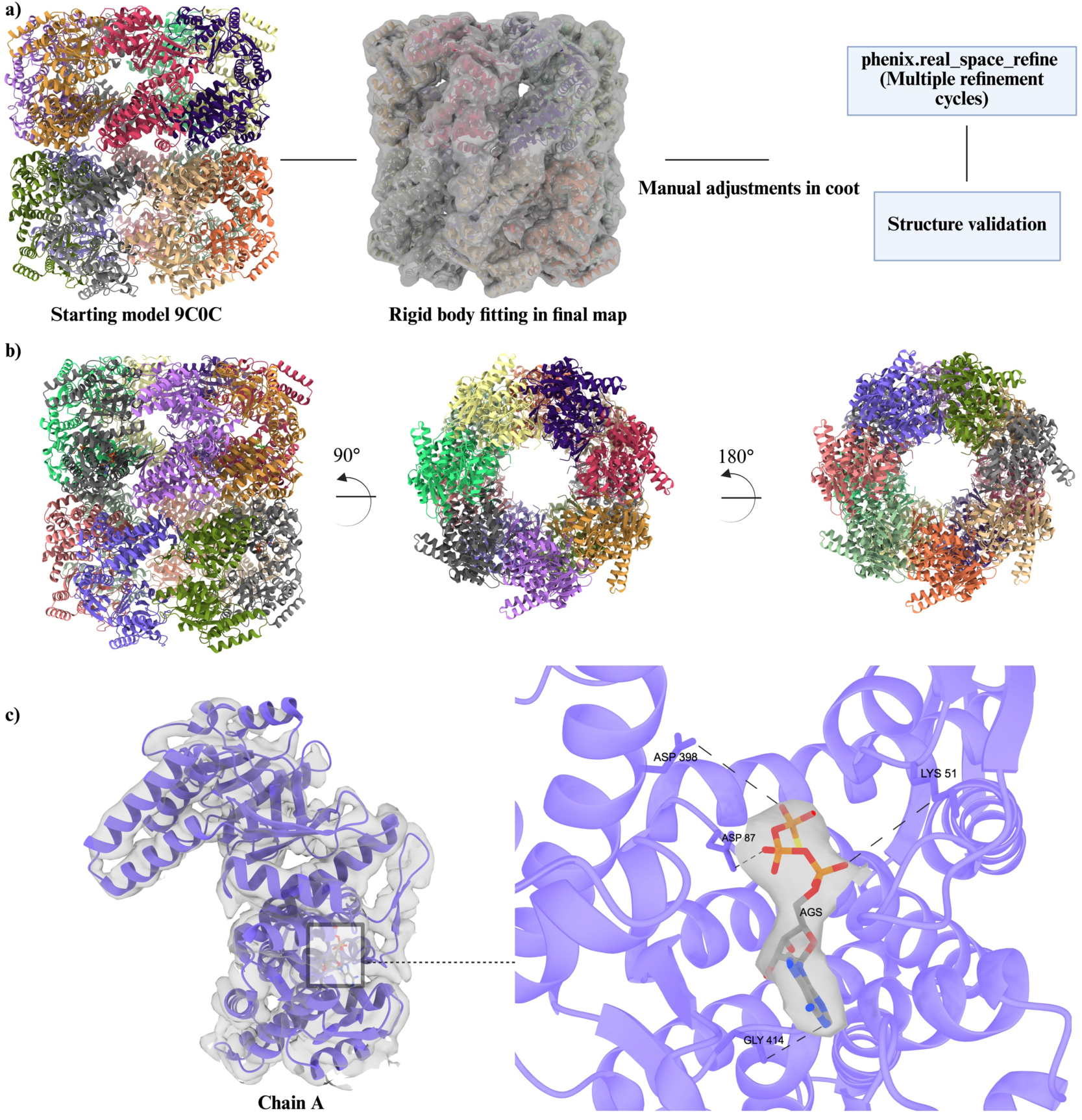
Model building and structural details of the GroEL-γATP bound complex. a) Schematic workflow of the model building and refinement process. The starting model for structural refinement was PDBID: 9C0C (GroEL-apo). Initially, the model was fitted in the final map using UCSF ChimeraX, followed by manual adjustments in COOT. Iterative refinement cycles were performed in Phenix (phenix. real_space_refine), and structure validation was carried out using MolProbity. The resulting models have favorable stereochemical parameters, including minimal deviation from ideal bond lengths and angles, and a low number of macromolecular backbone outliers. b) A side view of the multimeric GroEL complex is shown on the left, with each of the 14 subunits colored differently. The middle and right panels show top and bottom views of the complex respectively, to highlight the overall architecture and subunit organization. c) The left panel displays subunit A of GroEL bound to γATP (purple), with a boxed region indicating the ATPase active site. The right panel highlights this region in detail, showing the molecular surface enveloping the bound γATP. Key residues involved in ATP hydrolysis are labeled including: Lys51 (involved in ATP phosphate binding), Asp87 and Asp97 (coordinate the essential Mg²⁺ ion and help position water for nucleophilic attack), Asp398 (contributes to allosteric signaling and conformational changes), and Gly414 (provides structural flexibility for proper positioning of catalytic residues).

#### Structure Validation

During model validation Phenix does give a wide range of statistics for validation, as MolProbity is incorporated into the software (Davis *et al*., 2007). However, the model can also be validated in the MolProbity web server at https://molprobity.biochem.duke.edu/. An overview of data collection and validation statistics of all three structures are shown in **Table 6**. Here we will go through the validation procedure used for the GroEL-γATP complex. During model refinement in Phenix (Adams *et al*., 2010), global refinement statistics were monitored after each cycle to assess model quality and convergence. Upon completion of the refinement process, the final model-to-map cross-correlation coefficient (CCC) was 0.83, indicating a strong agreement between the atomic model and the experimental cryo-EM map. For geometry restraints, root mean square deviation (RMSD) for bonds and angles, quantifies how much the model’s geometry deviates from established ideal values. The low RMSD values for bonds (0.003 Å) and angles (0.599°) demonstrated that the model’s geometry was close to the expected ideal values. In addition, Z-scores (0.223 and 0.390, respectively) further confirm this, as they are very low and well within an acceptable range.

**Table 6.** Details of structural refinement of all three GroEL structures/models.

|  |  |  |  |
| --- | --- | --- | --- |
| **Data collection and processing** | **GroEL-ATP**<br>**PDBID: 9YKC**<br>**EMD-73044** | **GroEL-ADP**<br>**PMID: 9YNJ**<br>**EMD-73200** | **GroEL-apo**<br>**PDBID: 9YKE**<br>**EMD-73045** |
| **Magnification** | 81000 | 81000 | 81000 |
| **Voltage (kV)** | 300 | 300 | 300 |
| **Electron exposure (e-/Å<sup>2</sup>)** | 64 | 64 | 52.5 |
| **Defocus range (µm)** | -0.8 µm to -2.5 µm | -0.8 µm to -2.5 µm | -0.8 µm to -2.5 µm |
| **Pixel size** | 1.058 | 1.069 | 1.069 |
| **Symmetry imposed** | D7 | D7 | D7 |
| **Final map particles** | 261,106 | 78,179 | 42,776 |
| **Final map resolution** | 3.3 Å | 4.2 Å | 3.7 Å |
| **FSC threshold** | 0.143 | 0.143 | 0.143 |
| **Refinement details** |  |  |  |
| **Model used (PDB code)** | 8BL7 | 9C0C | 9C0C |
| **Resolution of starting model** | 4.40 Å | 3.41 Å | 3.41 Å |
| **Correlation Coefficient (cc_mask)** | 0.83 | 0.87 | 0.87 |
| **Map sharpening B factor (Å<sup>2</sup>)** | -60 | -100 | -40 |
| **MolProbity score** | 1.30 | 1.38 | 1.29 |
| **Clash score** | 3.1 | 4.0 | 3.0 |
| **Poor rotamers** | 0.1 % | 0 % | 0 % |
| **Ramachandran plot** |  |  |  |
| **Favored (%)** | 97.4% | 98.0% | 97.2% |
| **Allowed (%)** | 2.6% | 2.0 | 2.8% |
| **Disallowed (%)** | 0% | 0% | 0% |
| **CaBLAM outliers** | 2.3% | 1.4% | 1.6% |
| **Q-score** | 0.443 | 0.305 | 0.396 |

The all-atom clashscore for the GroEL-γATP was 3.1, which indicated very few steric clashes between atoms. The Ramachandran plot is a graphical way to visualize the energetically favored and allowed backbone conformations of amino acid residues in a protein. It plots the two main dihedral angles of the protein backbone, phi (ϕ) and psi (ψ). The Ramachandran plot had 0.00% outliers and a very high percentage of residues in the “favored” region (97.4% in this case). The rotamer outliers are amino acid side-chain conformations that are found in energetically unfavorable positions. Their presence could indicate potential errors in the model, as side chains typically adopt a limited set of stable conformations known as rotamers to minimize steric clashes. There were 0.1% rotamer outliers, which indicated favorable side chain conformations.

After reviewing the validation statistics, we checked each residue of our model to ensure its quality and adherence to stereochemical principles. This process helps identify potential errors or issues that could lead to future validation problems. Afterwards, we started the PDB validation, for this purpose the refined model (GroEL-γATP) and final map were uploaded to https://validate-rcsb-2.wwpdb.org/. The PDB validation report provides an overall summary, using color-coded flags to highlight potential issues. A green flag indicates a metric is within expected ranges; yellow suggests a potential issue, and red flags a significant problem. Another metric in the validation report was the Q-score (Pintilie *et al*., 2020), which quantifies the resolvability of individual atoms in a 3D density map. It measures how well the map values around a specific atom’s position correlate with an ideal, well-resolved Gaussian-like function, with a score of 1 indicating a perfect fit and a value closer to 0 or negative indicating poor resolvability or a poor fit, for all 14 chains in the our GroEL-γATP model, the Q-score was 0.443.

#### Structure deposition and repositories for cryo-EM data

Best practices for cryo-EM data deposition emphasize transparency, reproducibility, and community access. Final atomic models should be deposited in the Protein Data Bank (PDB), while corresponding 3D density maps must be submitted to the Electron Microscopy Data Bank (EMDB). For raw image data, deposition in the Electron Microscopy Public Image Archive (EMPIAR) is strongly encouraged, as it enables method development, validation, and training of new algorithms. These repositories provide persistent identifiers and standardized validation reports, ensuring compliance with funding, journal, and community guidelines. Using the OneDep system, models and maps can be deposited to the PDB and EMDB simultaneously through a single deposition workflow, streamlining the process and ensuring consistency across related datasets. Detailed instructions for deposition, including file formats and validation requirements, are available on the respective websites:

- **PDB:** https://www.wwpdb.org/
- **EMDB:** https://www.ebi.ac.uk/emdb/
- **EMPIAR:** https://www.ebi.ac.uk/empiar/

Following these practices aligns with the NIH 2023 Data Management & Sharing Policy and the NSF Public Access and Data Management & Sharing Plan requirements, promoting timely public access to publications and supporting data, and strengthening the transparency and reproducibility of federally funded research. It also supports the broader structural biology community in benchmarking and advancing the field.

### 7. Conclusions and Future Directions

Our transition from X-ray crystallography to single-particle cryo-EM provided critical insights into the structural characterization of ATAD2B, a large AAA+ ATPase and bromodomain-containing protein involved in chromatin regulation. This journey highlighted both the transformative potential of cryo-EM and the practical challenges associated with adopting this technique, including sample heterogeneity, co-purification of chaperones such as GroEL, and the steep learning curve for data processing and computational infrastructure. Despite these obstacles, we successfully established a workflow that integrates biochemical optimization, advanced vitrification strategies, and state-of-the-art image processing tools.

Our experience with ATAD2B highlights a critical question for cryo-EM practitioners: how much contamination can be tolerated before a targeted reconstruction becomes impractical? The contamination threshold is variable and can sometimes be overcome by collecting enough data, even if your particles are scarce (Ho *et al*., 2018). However, our experience suggests that when contaminant particles outnumber the target by an order of magnitude or more, as in our initial dataset where GroEL particles exceeded ATAD2B particles by nearly tenfold, the likelihood of achieving a high-resolution reconstruction of the target diminishes substantially. In such cases, even aggressive classification and filtering strategies may fail to recover sufficient particles for near-atomic resolution. Ultimately, when contaminant particles comprise more than ∼70–80% of the dataset, the time and computational cost required to isolate enough target particles becomes prohibitive, and a return to sample optimization is warranted. We chose to go back and improve the quality of the ATAD2B protein sample because 300 kV Krios data collection time is precious and our access was limiting. We also planned to continue structural studies on the ATAD2B AAA+ ATPase complex in the long-term, so it made sense to move to a different expression system to eliminate the contaminant. In other scenarios, a brute-force data collection strategy may make sense, particularly if microscope time and storage resources are abundant. This decision depends on several factors including, how much microscope time you have, what resolution you need in specific regions of the protein, and the intrinsic properties of your macromolecular complex. Highly symmetric contaminants, such as GroEL with D7 symmetry, refine rapidly and dominate classification, making it difficult to isolate minority species (Scheres, 2016). Highly dynamic complexes, flexible proteins, and heterogeneous samples will require more particles to reach high resolution, further increasing the impact of contamination. While dataset size and computational resources can partially offset these challenges, their success depends on practical constraints. Importantly, these factors can be manipulated, and adjusting the biochemical strategies remain the most effective approach. In our case switching expression systems from *E. coli* to Sf9 insect cells eliminated GroEL contamination and enabled collection of a homogeneous dataset suitable for high-resolution reconstructions. Additional troubleshooting approaches include editing the expression construct, altering the purification strategy, varying the buffer components and optimizing grid preparation parameters by testing alternative grid types, or employing additives such as detergents or graphene oxide supports to improve particle behavior (An *et al*., 2025). If cryo-EM grids exhibit poor particle distribution or preferred orientation adjusting the blotting conditions and plunge freezing strategy may be useful (e.g. Vitrobot versus Chameleon) (Levitz *et al*., 2022). An advantage in cryo-EM is that contaminants and heterogeneity can also be reduced computationally by employing selective particle-picking workflows (including neural network-based tools like Topaz) (Bepler *et al*., 2019). Topaz uses machine learning to selectively pick particles that match the target protein, reducing sample heterogeneity by excluding contaminants and junk particles. CryoDRGN and RECOVAR are additional deep-learning approaches that can addresses conformational heterogeneity by modeling continuous structural variability and conformational landscapes (Zhong *et al*., 2021). Despite these computational breakthroughs, structure determination by cryo-EM is still an iterative process and typically requires collection of multiple datasets (often three to four per structure) to achieve the desired resolution for a new structural target. In our case, improving the ATAD2B sample was the most efficient path forward, given our long-term goals and limited access to high-end instrumentation.

Looking forward, cryo-EM is poised to redefine structural biology by enabling visualization of macromolecular assemblies in unprecedented detail and in increasingly native contexts. Advances in detector technology, sample preparation, and computational methods—including machine learning-driven particle picking, heterogeneity analysis, and integrative modeling—will accelerate data processing and interpretation. Furthermore, the expansion of cryo-electron tomography and correlative workflows will bridge the gap between isolated complexes and cellular environments, offering a holistic view of molecular machines in action.

Together, these developments position cryo-EM to deliver not only higher resolution but higher information content: dynamic ensembles, contextualized structures, and integrative models aligned with cellular function. For the crystallographic community, the opportunity is clear—leveraging cryo-EM alongside crystallography and complementary biophysical approaches will deepen mechanistic understanding across scales, inform structure-guided discovery, and establish robust, community-vetted standards for the next decade of structural biology.

## Accession numbers

GroEL-gamma-ATP PDBID: 9YKC and EMDB: EMD-73044, GroEL-ADP PDBID: 9YNJ and EMDB: EMD-73200, and GroEL apoenzyme PDBID: 9YKE and EMDB: EMD-73045.

## Supporting information

Mass Spec supplemental data

## Acknowledgements

We gratefully acknowledge the staff at the National Center for Cryo-EM Access and Training (NCCAT) at the New York Structural Biology Center for their exceptional support and guidance throughout our training in single particle cryo-electron microscopy. Their expertise, patience, and dedication were instrumental in helping us develop the skills necessary for high-quality sample preparation, data collection, and analysis. We especially thank them for their hands-on instruction and thoughtful mentorship during our visits, which greatly enriched our understanding and capabilities in cryo-EM. Special thanks to Christina Zimanyi and Eugene Chua for their critical feedback and thoughtful editing of this manuscript.

## Author Contributions

HZ, KLM, AKS, MAC, and KCG conceptualized the work. All authors contributed to the literature review and analysis of current research, and all authors drafted, revised, and approved the manuscript. All authors approved the submission and agreed to be responsible for their contributions.

