## Supplemental Information for "Breaking Barriers: Transitioning from X-ray Crystallography to Cryo-EM for Structural Studies"

**Table of Contents:**

**Supplemental Methods:** Expression and purification of the human ATAD2B protein from *E. coli*…………………………………………………………………………………...…...…….2

**Supplemental Figure S1:** Representative SDS-PAGE gel of purified ATAD2B and excised bands for proteomic analysis…………………………………………………………………...4

**Supplemental Figure S2:** Mass spectrometry identification and quantification of purified ATAD2B sample components (provided as a separate file ZafarH_Mass_Spec_supplemental_data.xls)

### Supplemental Methods:

*Expression and purification of the human ATAD2B protein from E. coli*

Human ATAD2B (residues 380-1458, UniProt ID: Q9ULI0) was cloned into a pGEX-6P-1 plasmid containing an N-terminal GST-tag, followed by a PreScission Protease Cleavage site and the ATAD2B sequence that had been codon optimized and synthesized by GenScript (Piscataway, NJ, USA). The plasmid was transformed into *Escherichia coli (E. coli)* Rosetta 2(DE3)pLysS competent cells (Novagen). A 4 L culture of *E. coli* cells containing the GST-tagged ATAD2B protein was grown in Terrific broth (TB) shaking at 225 rev/min in 37 °C until they reached an OD600 of 1.0, and then the temperature was dropped to 20 °C for 1 h before addition of 0.1 mM isopropyl β-D-1-thiogalactopyranoside (IPTG). The cells were grown for an additional 16 h at 20 °C and harvested by pelleting the cells using centrifugation at 10,000 x g for 20 min. The harvested bacterial pellet was resuspended in 100 mL of lysis buffer (50 mM Tris, pH 7.5, 500 mM NaCl, 5% glycerol, 2 mM $MgCl_2$, 1 mM dithiothreitol (DTT)) supplemented with 100 mM PMSF, 1 protease inhibitor tablet (Pierce protease inhibitor tablets, EDTA-free, Thermo Fisher), and lysozyme (1 mL). Cells were disrupted by sonication, and the resulting lysate was clarified by centrifugation at 10,000 RPM for 20 min. The supernatant was incubated with 10 mL of glutathione agarose resin (Thermo Scientific; binding capacity ~40 mg GST-tagged protein per mL) for 2 h at 4 °C with gentle agitation. After binding, the resin was collected by centrifugation (500 × g, 5 min) and transferred to a 25 mL Econo-Column (Bio-Rad). The column was washed with four volumes of wash buffer (25 mM Tris, pH 7.5, 500 mM NaCl, 5% glycerol, 2 mM $MgCl_2$, 1 mM DTT). GST-tagged ATAD2B protein was eluted off the beads with 20 mM reduced glutathione, and dialyzed into wash buffer to remove the reduced glutathione overnight. The protein was concentrated to ~500 μL using an Amicon® Ultra

Centrifugal Filter with a 10-30 kDa molecular weight cut-off for further purification using size-exclusion chromatography (SEC). The ATAD2B protein was applied to a Superdex 200 Increase 10/300 GL column on an ÄktaPrime system (GE/Cytiva) in wash buffer and the peak fractions corresponding to the ATAD2B construct (see Figure 2C, peak at 7-10 mL) were pooled and concentrated to a final concentration of 2.0-2.5 mg/mL. Protein purity was confirmed by sodium dodecyl sulfate-polyacrylamide gel electrophoresis (SDS-PAGE) gels stained with Bio-Safe Coomassie G-250 Stain (Bio-Rad). Protein concentrations were determined with NanoDrop (Thermo Scientific, Waltham, MA, USA). For structural studies the ATAD2B protein was dialyzed into cryo-EM buffer (20 mM HEPES pH 7.5, 150 mM NaCl, 5% glycerol, 2 mM $MgCl_2$, 1 mM DTT) and flash frozen in liquid nitrogen and stored at -80 °C for future experiments.

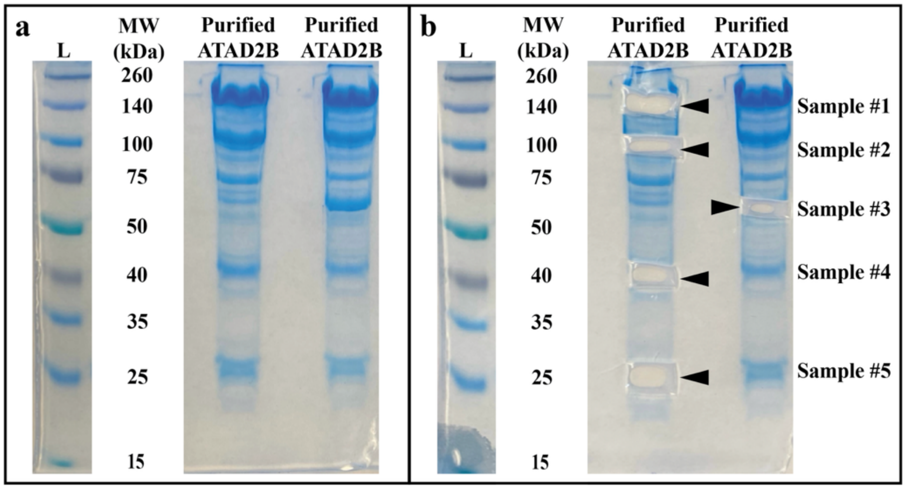


**Supplemental Figure S1: Representative SDS-PAGE gel of purified ATAD2B and excised bands for proteomic analysis.** The ATAD2B protein was expressed and purified from *E. coli* as described in the methods section. a) After size-exclusion chromatography the final pooled and concentrated ATAD2B sample in wash buffer (25 mM Tris pH 7.5, 500 mM NaCl, 5% glycerol, 2 mM $MgCl_2$, 1 mM DTT) was run on a freshly cast 10% SDS-PAGE gel and stained with Coomassie blue. L denotes the molecular weight (MW) standards lane with the MW for each band labeled followed by two lanes of the same purified ATAD2B sample. b) Five protein bands (Samples #1-5) were cut from the gel lanes using a fresh razorblade for each band. The bands chosen for mass spectrometry analysis correspond to bands with molecular weights of 140 kDa, 100 kDa, 70 kDa, 40 kDa, and 25 kDa. Each band was directly placed in an autoclaved Eppendorf ube containing ~500 µL of Milli-Q $H_2O$, sealed with parafilm, and shipped overnight to the Mass Spectrometry Proteomics Shared Resource Facility at University of Colorado-Anschutz. A liquid sample (Sample #6) of the purified ATAD2B protein that was run on the gel was also included in this shipment. Mass spectrometry data were analyzed using the Mascot Server (https://www.matrixscience.com/server.html) to search the human protein database and ATAD2B was identified in all five samples. Later, after opening the search to both human and *E. coli* protein databases, GroEL was also identified in Samples 2 and 3. For full list of identified proteins and relative abundances see the additional supplementary file ZafarH_Mass_Spec_supplemental_data.xls.
